# NGN3 oscillatory expression controls the timing of human pancreatic endocrine differentiation

**DOI:** 10.1101/2024.01.10.574974

**Authors:** Anzy Miller, Veronica Biga, Andrew Rowntree, Mariya Chhatriwala, Florence Woods, Elli Marinopoulou, Anoushka Kamath, Ludovic Vallier, Nancy Papalopulu

## Abstract

Understanding protein expression dynamics is crucial for the mechanistic understanding of cell differentiation. We investigate the dynamics of NGN3, a transcription factor critical for pancreatic endocrine development, including their function and decoding mechanisms. A knock-in endogenous reporter shows that the expression of NGN3 protein oscillates with a 13-hour periodicity in human iPS-derived endocrine progenitors and is switched off as cells differentiate to β-like and α-like cells. Increasing the stability of NGN3 protein results in one broad peak of expression instead of oscillations, with a larger peak to trough fold-change. This leads to precocious endocrine differentiation to both β-like and α-like cells and precocious expression of key NGN3 target genes. Single-cell analysis of dynamics, mathematical modelling and bioinformatics suggest that decoding of NGN3 oscillations occurs by fold-change detection via an incoherent feedforward motif that explains both normal and precocious differentiation. Our findings suggest that oscillatory NGN3 dynamics control the timing of differentiation, but not fate specification.

## Introduction

The pancreatic endocrine islets contain the cells that are central for glucose homeostasis: INSULIN (β)-, GLUCAGON (α)-, SOMATOSTATIN (δ)-, PANCREATIC POLYPEPTIDE(γ)- and GHRELIN(ε)-secreting cells. Diabetes results from the inability of these pancreatic cells to manage blood glucose levels, caused by either the autoimmune destruction of β-cells (Type 1 diabetes) or β-cell dysfunction paired with insulin resistant tissues (Type 2 diabetes).

In the mouse, all pancreatic islet cells originate from a NEUROGENIN3 (NGN3, encoded by *Neurog3* gene)-expressing pancreatic endocrine progenitor and loss of NGN3 results in no islet cells produced^1,2^. In humans, mutations in *NEUROG3* have been identified in patients who develop diabetes^3–5^ and loss of NGN3 either results in very few, or a complete loss of islet cells highlighting NGN3 is also very important for human pancreatic cell development^6–10^.

NGN3 is a basic helix-loop-helix transcription factor, known to directly activate a broad range of genes critical for islet cell development in mice and humans^9–21^. While genome wide approaches have helped us understand the gene regulatory networks (GRNs) that occur downstream of NGN3^10,22,23^, it is still unknown how NGN3 impacts on the activation of the different endocrine GRNs, and how these GRNs are activated at the correct time.

Recent single-cell live imaging has shown that several proteins, rather than being in either a steady “on” or “off” state, can be expressed in a highly dynamic way, for example oscillate with a periodicity of a few hours (P53, HES family, ASCL1, DLL1). These dynamics have been shown to be functionally important developmentally, for neurogenesis^24–27^, somitogenesis^28–31^ and myogenesis^32^. Recently, oscillatory dynamics of HES1 and DLL1 were also observed in pancreas development, and found to be important for the proliferation of pancreatic progenitors^33^, implying the likelihood that these dynamics are widespread throughout development. These studies highlight the importance of monitoring transcription factor protein dynamics at single cell level in order to understand the regulation of cell state transitions.

In pancreas development, the upstream repressor and downstream target of NGN3 (HES1 and DLL1 respectively) are expressed in an oscillatory manner^33^. The possibility that NGN3 has dynamic expression is important to investigate since NGN3’s level, stability and timing of expression have been shown to be important for endocrine differentiation^34–36^. It is possible that NGN3 dynamics could affect cell fate, since evidence suggests that NGN3 positive cells are post-mitotic^37^ and each NGN3+ cell gives rise to a single type of islet cell, indicating that the fate decision could be finalised within the NGN3 expressing window^38^. In addition, NGN3 dynamics could also affect differentiation timing, since human endocrine cell differentiation is spread out across at least a 3-month period^39–41^, but the regulatory mechanisms that control the timing of differentiation are unknown.

Although many signalling factors have oscillatory dynamics ^42,43^, little is known about **how** the dynamics of transcription factors are decoded by the cell, and the effect this has on differentiation. Evidence from various signalling pathways suggest that downstream factors can respond to changes in frequency, fold-change and the phase of input signals. For example, the oscillation frequency of TNFα influences the ability of NFκB to translocate into the nucleus and regulate transcription^44^, and in the WNT pathway, β-CATENIN is responsive to the fold-changes of WNT signal rather than the absolute levels^45^. Thus, to understand how dynamic transcription factors regulate cell fate transitions, we need to consider not only protein abundance but also duration, frequency and fold change of these factors^43,46^.

In this work, we focus on the protein expression dynamics of NGN3 which is currently overlooked, and whether those dynamics play a role in cell-fate decisions and/or in the timing of differentiation, and how the dynamics are decoded by the cell. We created an endogenous homozygous NGN3::mVenus fusion protein in human induced pluripotent stem (iPS) cells and by utilising a human pluripotent stem cell (hPSC) differentiation protocol, we provide a detailed characterisation of human NGN3 expression in pancreatic endocrine progenitors at the single cell level. We report that NGN3 protein oscillates during its window of expression, and that these oscillatory dynamics do not correlate with cell fate specification since they are observed with the same characteristics irrespective of whether they give rise to β-like or α-like cells. However, by generating a more stable NGN3 protein (by creating a phosphomutant-NGN3), we change the oscillatory dynamics and show that, instead, NGN3 can be predominately expressed as one long pulse. This resulted in precocious differentiation compared to WT cells but did not bias differentiation towards a particular cell type. We found evidence to suggest that the fold-change of NGN3 oscillations is decoded via an incoherent feed-forward gene regulatory motif and that oscillatory expression of the master regulator NGN3 does not control fate specification, but instead is required to distribute differentiation over time in the human pancreatic endocrine.

## Results

### NGN3 oscillates in human endocrine progenitors

Directed differentiation of human induced pluripotent stem cells (hiPSCs) into pancreatic cells was undertaken using a well-controlled hiPSC pancreatic differentiation model^47^ (Fig.1A). An essential step in the transition to endocrine cell specification is represented by endocrine progenitors which express the transcription factor NGN3^8^. As we are interested in understanding the dynamic behaviour of NGN3 during the progenitor to endocrine transition, we generated endogenous knock-in hiPSC cell lines by fusing the fluorescent protein mVenus (including 3XFLAG, HA and linker sequences) in-frame to the 3’ end of *NEUROG3* (just upstream of the stop codon) using CRISPR/Cas9 homology directed repair (HDR) (Fig.1B). Single cell clones were verified by sequencing of both *NEUROG3* alleles. The addition of mVenus to NGN3 created a faithful reporter of NGN3 (Fig.S1A), and did not impact on the function of the protein since homozygous NGN3::mVenus hiPS cells were able to differentiate into immature endocrine cells, expressing hormones which mark their fate (INSULIN, using C-PEPTIDE (CPEP) as a marker; GLUCAGON, GCG and SOMATOSTAIN, SST) (Fig.1C).

**Figure 1.**
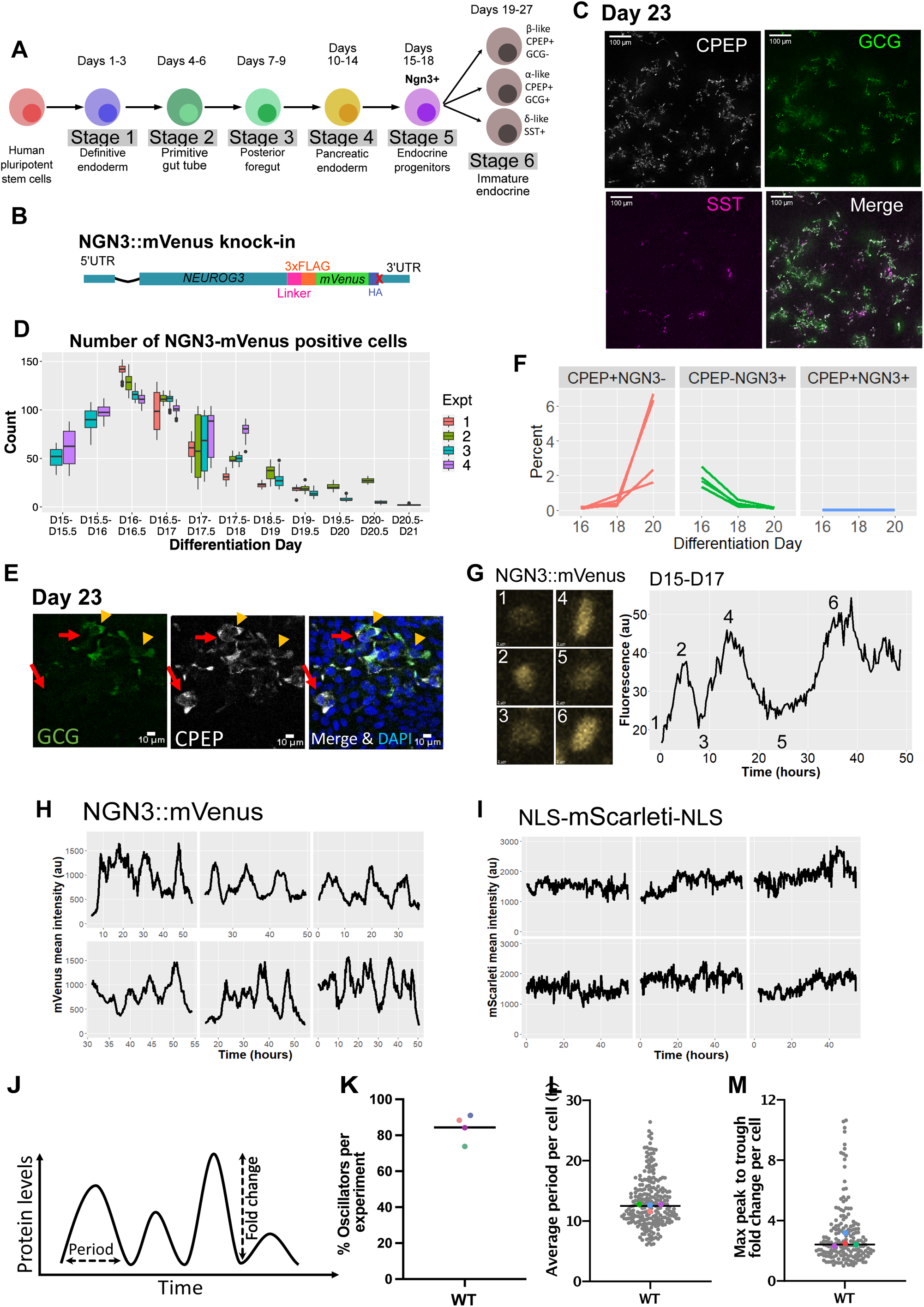
Timelapse monitoring of endogenous human NGN3 at single-cell level using an in-vitro pancreas differentiation protocol. **(A)** An overview of the in-vitro differentiation protocol from human pluripotent stem cells to produce endocrine progenitors which express NGN3 (day 15-18) and further differentiated endocrine cell types (from day 19 onwards) identified with C-PEPTIDE (CPEP), GLUCAGON (GCG) and SOMATOSTATIN (SST). **(B)** Schematic of the endogenous fusion knock-in designed for visualising NGN3 by direct fusion to fluorescent protein mVenus (see also Materials and Methods). **(C)** Single-cell expression of endocrine markers CPEP, GCG and SST observed by immunofluorescence at day 23 of protocol (A). Scale bar is 100µm. **(D)** Quantification of absolute NGN3::mVenus+ cell numbers observed during pancreas differentiation days 15 to 21 across 4 independent experiments. The number of positive cells were counted per time point of the timeseries per field of view, and then binned into 12hr bins. Boxes indicate median and interquartile range; dots indicate outliers. **(E)** At differentiation day 23, cells with single expression CPEP+/GCG-(red arrows) can be observed alongside cells with double expression CPEP+/GCG+ (orange arrowheads). Scale bar is 10µm. **(F)** Percentage of C-PEPTIDE and NGN3 expressing cells determined by flow cytometry during differentiation days 16 to 20; lines indicate 4 independent experiments. Example flow plot shown in Fig. S1D. **(G)** Representative examples of a NGN3::mVenus expressing cell observed over time (left panel) showing oscillations in average fluorescent intensity (right panel); corresponding variations in nuclear marker NLS-mScarleti in the same cell shown in Fig. S1E. **(H-I)** Additional examples of NGN3::mVenus **(H)** and nuclear marker NLS-mScarleti-NLS **(I)** intensities observed in different cells over time. **(J)** Schematic illustrating the dynamic parameters measured from timeseries data. **(K)** Percent of oscillators expressing NGN3::mVenus observed in 4 independent experiments determined through a false discovery rate statistical test (Materials and Methods); colour dots indicate experiments; line indicates median; sample size 287 cells overall. **(L)** Estimation of period from oscillatory fractions in **(K)** determined through a Gaussian Process approach (Materials and Methods); colour dots indicate median of independent experiments; gray markers indicate single-cell tracks; line indicates median overall. **(M)** Estimation of maximum variations observed between a trough and a peak reported as a peak to trough fold change (Materials and Methods); gray markers indicate cells; colour dots indicate median per experiment; line indicates median overall.

At the population level, the window of expression of NGN3 is transient, consistent with previous data from in vitro differentiations^48–50^. The intensity and number of endocrine progenitor cells expressing NGN3::mVenus peaks at day 16 and then declines until day 19 when very few cells express NGN3 (Fig.1D,S1B). NGN3::mVenus+ cells did not divide, consistent with the evidence that NGN3 expression causes exit from the cell cycle^37^. This pancreatic differentiation model mainly produces two immature endocrine cell types (Fig.1E): β-like cells only expressing INSULIN hormone (CPEP+GCG-), and α-like cells, expressing both INSULIN and GLUCAGON (CPEP+GCG+)^40,50–53^. Some delta-like cells, expressing the hormone SOMATOSTATIN (SST+) are also produced but very few in comparison to the other two types (Fig.1C, S1C). A small number of differentiated cells expressing INSULIN are first observed at day 18, which increases at day 20, and these CPEP+ cells do not co-express NGN3, showing NGN3 has turned off before differentiation occurs (Fig. 1F, S1D), in agreement with previous studies^54,55^.

Using time-lapse imaging of NGN3::mVenus expression at single cell level between days 15 and 19, we observed that NGN3::mVenus expression shows repeated peaks and throughs in endocrine progenitors (Fig.1G,H) suggestive of oscillatory expression. As expected, the nuclear marker NLS-mScarleti-NLS, stably expressed from a constitutive promoter, was steadily expressed with only minor variations over time (Fig.1I, S1E). Using the FisherG statistic, which is a measure of dynamic activity at single cell level, we show that NGN3::mVenus fluorescence is significantly more dynamic compared to NLS-mScarleti-NLS fluorescence and both are above fluctuation levels observed in cells not expressing mVenus or mScarleti (representative of detector noise, Venus-null and Scarlet-null respectively) (Fig.S1F).

To find out whether the dynamic expression was oscillatory, we analysed single-cell timeseries (tracks) using customisable Gaussian Processes pipeline previously described^24,56^. We found that 85-90% of cells had oscillatory NGN3::mVenus expression (Fig.1J, S1G, S1I), and none of the NLS-mScarleti-NLS timeseries passed the statistical test due to a significantly reduced log-likelihood ratio (LLR) statistic compared to NGN3::mVenus cell timeseries (Fig.S1G). In addition, we found no significant differences in the percent of oscillatory cells between homozygous and heterozygous NGN3::mVenus cells lines (Fig.S1H,I). These results were unaffected by differences in sample size and track length (Table S4).

To quantify further the NGN3 dynamics, we analysed the average period per cell (the amount of time it takes for one complete oscillation, e.g. from peak to peak of the oscillator) and the maximum fold change per cell (the ratio between peak and trough-see Fig. 1J for an illustration of these terms). Homozygous and heterozygous NGN3::mVenus cells oscillated with a median period of 13 and 11 hours respectively, broadly reproducible between experiments (Fig.1K, S1J), and the median of the maximum peak to trough fold change per cell was 2.4x (Fig. 1L). The presence of widespread oscillations in both heterozygous and homozygous conditions (Fig. S1I) indicated that NGN3 oscillates in-phase between alleles, and in independent knock-in cell lines. We proceeded to use the homozygous NGN3::mVenus cells (accounting for total NGN3) for further analysis.

Overall, our results show that the majority of human endocrine progenitors exhibit a transient time window (across 4-5 days) of oscillatory NGN3 protein expression with a period of 13 hours and maximum fold change of 2.4x, and expression is terminated before cells terminally differentiate.

### There is no difference in NGN3 expression between cells that differentiate into β-like or α-like cells

To understand the role of NGN3’s oscillations on endocrine cell fate decisions, we first investigated whether there were any differences in NGN3’s expression pattern between cells that go on to become β-like or α-like cells. As mentioned previously, the two main differentiated cells produced in this *in vitro* differentiation protocol express either C-PEPTIDE (a marker for INSULIN) alone (β-like cells), or C-PEPTIDE and GLUCAGON (α-like cells) and as such we explored the differences between these two cell fate outcomes. We performed a time-lapse movie of the endocrine progenitor cells expressing NGN3, and then using the nuclear marker NLS-mScarleti-NLS, continued to track these cells after NGN3 had turned off. At Day 19 or Day 20, these differentiations were then fixed and immunofluorescence was performed using antibodies against C-PEPTIDE and GLUCAGON (Fig. 2A). These experiments were performed on gridded dishes to be able to locate the same cells between time-lapse microscopy and immunofluorescence staining. Using this method, across three independent experiments, we were able to track NGN3’s dynamics in 124 cells from which 97 cells became β-like cells and 27 cells became α-like cells by Day 19 or 20.

**Figure 2.**
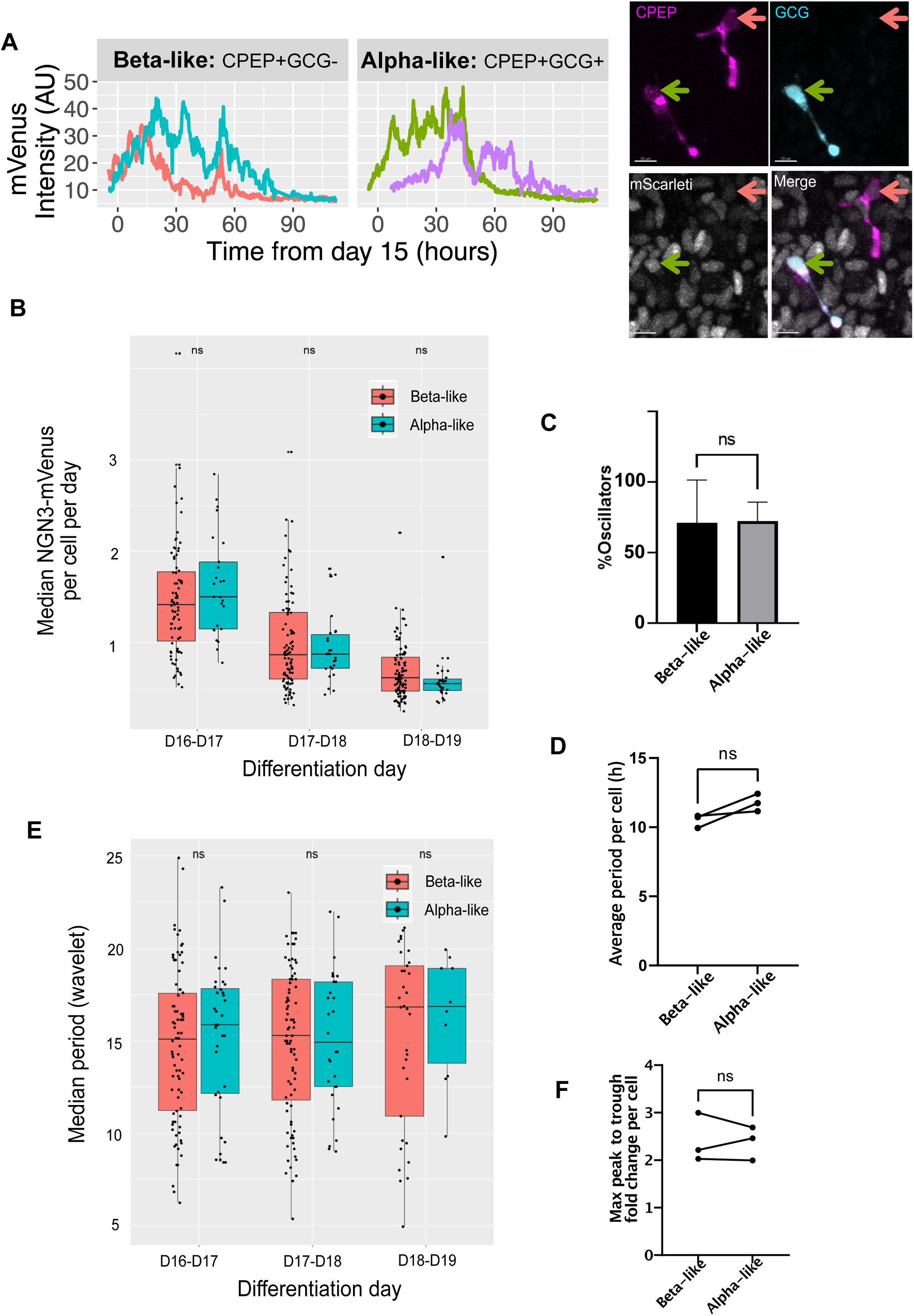
Oscillatory dynamics of NGN3::mVenus in differentiating progenitor cells. **(A)** Representative examples of NGN3 expression observed at single-cell level in cells that become CPEP+/GCG-(left panel) and CPEP+/GCG+ (middle panel) as determined by timelapse imaging and immunofluorescence at the end of the timelapse (right panel), see Materials and Methods-fate mapping. Colour arrows correspond to timeseries of the same colour in the left and middle panel. **(B)** Quantification of normalised median NGN3::mVenus intensities observed per cell in cells that become Beta-like cells (CPEP+/GCG-) versus Alpha-like cells (CPEP+/GCG+); points indicate cells, binning 24h, boxes indicate median and interquartile range from 3 independent experiments. **(C)** Percentage of oscillators observed in 3 independent experiments across the CPEP+/GCG- and CPEP+/GCG+ fractions of cells; bars indicate mean and SD per experiment, paired t-test, two tailed non-significant, p=0.9076. **(D)** Estimation of period (using a Gaussian Process approach-Materials and Methods) from oscillatory Beta and Alpha-like cells identified in (**C)**; markers indicate medians per experiment compared by a paired t-test, two tailed non-significant p=0.1129. **(E)** Estimation of period observed in a 12h time window (using Wavelet-Materials and Methods) observed in cells that become Beta and Alpha-like; points indicate cells; binning 24h; boxes indicate median and interquartile range from 3 independent experiments. **(F)** Maximum peak to trough fold change observed in CPEP+/GCG- and CPEP+/GCG+ fractions; markers indicate medians per experiment compared by a paired t-test, two tailed non-significant p=0.8593.

To investigate whether NGN3’s expression differs in cells that acquire different fates, we first assessed the levels over time of NGN3. We calculated the median mVenus intensity from each raw cell trace per 12 hours and plotted these values from Day 16 - Day 19. We found no significant differences in levels across the time-window of NGN3 expression between the cells that become α- or β-like cells (Fig.2B). Using the Gaussian Process-based statistical method described above, we then sought to analyse the dynamics of these cells. We found that NGN3 was equally likely to oscillate regardless of resulting fate (Fig.2C) and the percentage of oscillators across both fates were very similar to what we had found previously from the larger dataset, analysing all cells in the population (Fig.1J). We also found that there was no difference between the period of NGN3 oscillations in cells that acquire a different fate (Fig.2D). To account for any changes in period over time between cell types, we also compared their corresponding NGN3::mVenus traces using a Wavelet transforms platform (PyBoat) ^57^. This allows us to analyse periodicity over time and identify potential changes in the period that could be masked by averaging all periods across the trace. We calculated the median period per cell per 12 hours from Day 16 to Day 19 and found no significant differences in periods between NGN3::mVenus+ cells that go on to become α- or β-like cells (Fig.2E). In addition to no change in period (Fig.2D,E), we found no significant difference in the maximum peak to trough fold change (the median of each experiment being within the range of 2x to 3x) of the two cell groups (Fig. 2F) consistent with the wider population (Fig.1L).

In conclusion, we find no evidence to suggest that the expression of NGN3 is associated with cell fate specification of β-like or α-like cells, suggesting that neither oscillations nor levels are decoded in differential fate acquisition.

### Increasing the stability of NGN3 impairs oscillations

In order to understand the role of NGN3 oscillations, we created an endogenous homozygous phosphomutant NGN3::mVenus hIPS clonal cell line (named PM-NGN3::mVenus hereafter) where we mutated five serine-proline (SP) sites to alanine-proline (AP) at the C-terminus of the protein (Fig.3A). These mutations are known to increase the stability of the protein ^36,58^, which we also validated (Fig.S2A). We hypothesized that increasing the stability of NGN3 would impact on its oscillatory dynamics, as the generation of oscillations relies on many molecular requirements including protein instability ^59^. Indeed, in the majority of PM-NGN3+ hIPS-derived endocrine progenitors, the protein expression no longer oscillated over time, and instead the cells showed one long pulse of PM-NGN3 expression (Fig.3B, S2C). Interestingly, the population level window of PM-NGN3 expression was very similar to WT-NGN3 expression: cells started to express PM-NGN3 at day 15, with the highest number of cells expressing NGN3, with the highest intensity, at day 16 (Fig.3C, S2B). Just like WT-NGN3+, there are fewer PM-NGN3+ cells from day 17 onwards, until very few cells have detectable NGN3 expression by day 19 (Fig.3C, S2B).

**Figure 3.**
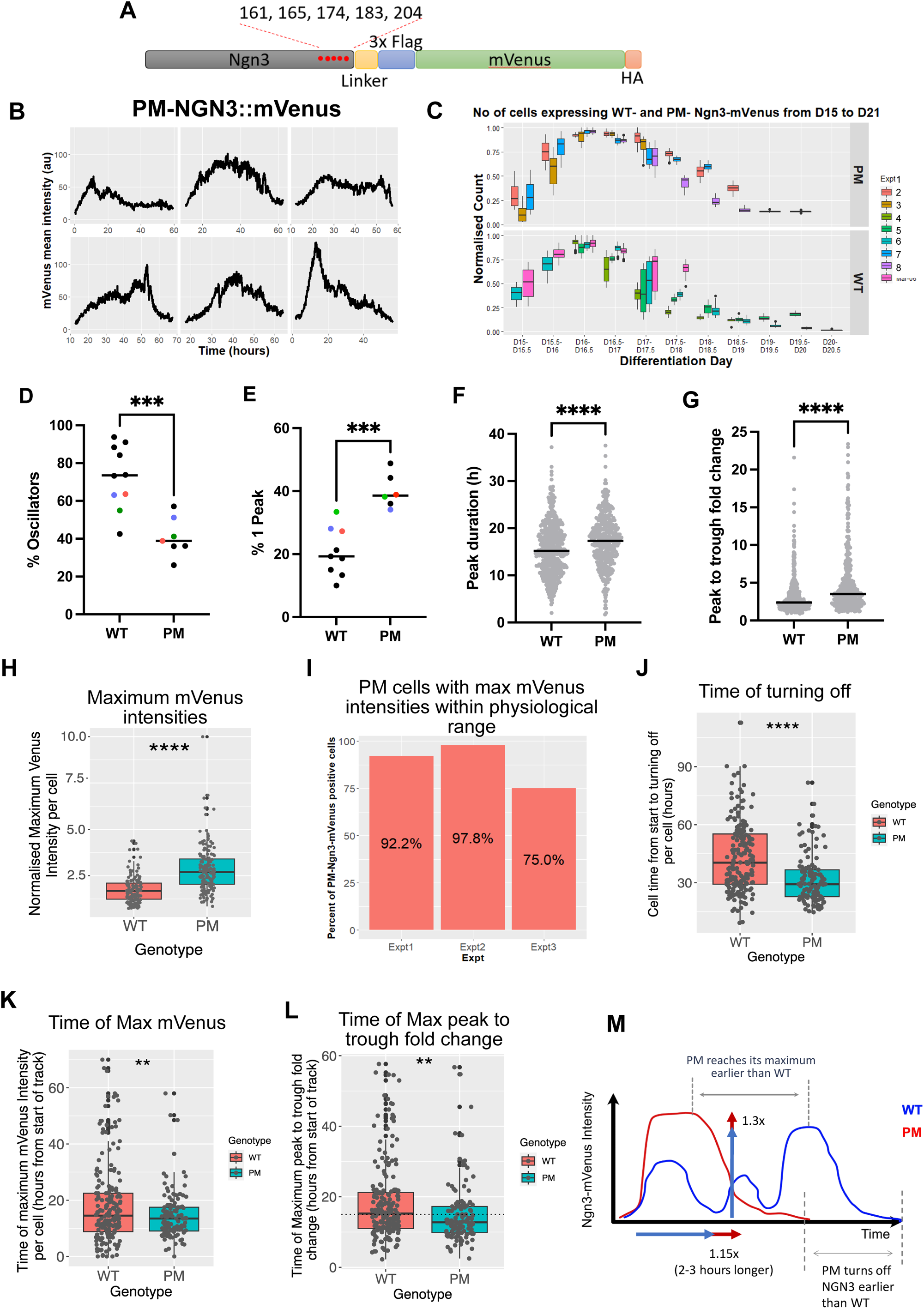
Comparative analysis of dynamics in a phosphomutant NGN3 versus wild-type. **(A)** Schematic of the phosphomutant (PM) knock-in where NGN3 is fused to mVenus. Red dots indicate the location of mutated SP sites in the C-terminus. **(B)** Representative examples of dynamic fluctuations observed in pancreatic progenitor cells expressing PM-NGN3::mVenus during differentiation days 15 to 19; intensity values reach high above background (see **Fig S2C**). **(C)** Fraction of cells expressing PM-NGN3::mVenus (top panel) and WT-NGN3::mVenus (bottom panel) monitored over differentiation days 15 to 21; cell counts were normalised to the WT and PM counts at day 16; binning 12h; bars indicate median and interquartile range of individual experiments; dots indicate outliers; 4 independent experiments per cell type. **(D)** Percentage of NGN3 oscillators determined by a false discovery rate test (Materials and Methods) observed in WT versus PM; markers indicate independent experiments with colour dots indicating co-culture experiments, lines indicates medians, statistical significance determined with a t-test, two-tailed p<0.001***. **(E)** Percentage of tracks with a single peak dynamic determined by auto-correlation function (ACF) analysis (Materials and Methods) observed in WT versus PM; markers indicate independent experiments with colour dots indicating co-culture experiments, lines indicate medians, statistical significance determined with a t-test two-tailed p<0.01**. **(F)** Individual peak durations determined by ACF (Materials and Methods) observed in WT versus PM cells; gray markers indicate single peaks, lines indicate medians, statistical significance determined with a Mann-Whitney test p<0.01**. **(G)** Estimation of maximum peak to trough observed in WT versus PM cells; gray markers indicates tracks, lines indicate medians, statistical significance determined with a Mann-Whitney test p<0.001***. **(H)** Quantification of maximum NGN3-mVenus intensity observed WT and PM cells from 3 independent co-culture differentiations, each experiment normalised to the maximum per experiment. Points indicate cells, boxes indicate median and interquartile range, statistical significance determined with a t-test p<0.0001****. **(I)** Percentage of cells with PM-NGN3::mVenus expression within the physiological range of WT-NGN3::mVenus level; bars indicate %PM cells with mVenus expression within the range of maximum WT mVenus intensities across 3 independent co-culture experiments. **(J)** Comparison of time duration until single WT versus PM cells switch off NGN3, expressed in hours from the start of NGN3 expression per cell, from 6 differentiations (3 cocultures, and 3 WT differentiations). Gray points indicate single cells, boxes indicate median and interquartile range, statistical significance determined with t-test p<0.0001****. **(K)** Time duration until single WT and PM cells reach maximum NGN3; time duration is calculated from start of NGN3 expression in each track; points indicate cells from 6 independent differentiations (3 cocultures, and 3 WT differentiations); boxes indicate median and interquartile range, statistical significance determined with a t-test p<0.01**. **(L)** Comparative analysis of time to reach maximum NGN3 peak to trough fold change per cell observed in WT versus PM; time is calculated from start of NGN3 expression in each track; markers indicate cells from 6 independent experiments (3 cocultures, and 3 WT differentiations), boxes indicate median and interquartile range, statistical significance determined with a t-test p<0.01**; dashed line indicates 15hours. 50% (WT) vs 38% (PM) of cells are above 15hours. **(M)** Diagram summarising the main differences in dynamic expression observed in WT versus PM.

We explored the dynamics of PM cells in independent differentiation experiments, but also to control for variation between differentiations, we set up co-culture differentiations of PM-NGN3::mVenus hiPS cells in the same well as WT-NGN3::mVenus hiPS cells. To achieve this, we made use of the NLS-mScarleti-NLS nuclear marker stably expressed in the WT cells (as mentioned above) which allowed us to distinguish between genotypes once mixed (Fig. S2D). For the co-cultures we mixed 50% WT-NGN3::mVenus hiPS with 50% PM-NGN3::mVenus hiPS cells at the beginning of the differentiation, and this 1:1 ratio was maintained throughout the differentiation (Fig.S2E). The percentage of NGN3::mVenus positive cells that were WT or PM was also 50% each at day 16 (Fig.S2F), which shows that changing the stability of NGN3 does not affect the number of cells that turn on NGN3 in this differentiation protocol.

Across culture conditions (co-culture and independent differentiations), we found a significant reduction in the percentage of oscillatory cells in endocrine progenitor cells expressing PM-NGN3 compared to WT-NGN3 with a median of only 39% of PM-NGN3 cells passing as oscillatory (Fig.3D, S2G). The number of cells analysed and the length of the single-cell tracks did not impact on this result (Table S4). We complemented this analysis by auto-correlation methods which also enabled us to distinguish a single pulse from periodic signals. We find there are significantly more PM-NGN3 endocrine progenitor cells that express only one peak of NGN3 (Fig. 3E, S2H), and that the median peak duration is elongated by 2¼ hours, in comparison to WT-NGN3 expressing endocrine progenitor cells (Fig.3F, S2I). In addition to peak elongation, the maximum peak to trough fold change is increased in PM-NGN3 compared to WT-NGN3 (Fig.3G, S2J), from a median of 2.4x to 3.5x overall.

To gain further insight into the differences between PM-NGN3- and WT-NGN3 dynamics, we analysed additional parameters of the dynamic expression. In mixed WT and PM conditions, the maximum NGN3::mVenus intensity level per cell of PM-NGN3+ cells was on average 1.3x higher than in WT-NGN3+ cells (Fig.3H), but interestingly the majority of PM-NGN3+ cells were within the same range of what WT-NGN3+ cells normally express (Fig.3I). Therefore, while WT-NGN3+ cells can express NGN3 at this higher level, they do so less often than PM-NGN3+ cells. We also found that PM-NGN3+ cells turn off NGN3 earlier than WT-NGN3+ cells at the single-cell level (Fig. 3J, S2K). We also observed that the time it takes for a PM-NGN3+ cell to reach its maximum NGN3::mVenus intensity level was significantly shorter than in WT-NGN3+ cells (Fig. 3K, S2L), and we find that 50% of WT cells require more than one peak (approx. 15hrs) to reach their largest amplitude peak, while 62% of PM achieve this in less than 15hours of starting to express NGN3 (Fig. 3L, S2M).

In conclusion, we find increasing the stability of NGN3 reduces the number of cells that have oscillatory NGN3 expression and increases the likelihood of one peak of expression with longer duration. In addition, PM-NGN3 cells have an earlier, higher fold change peak and individual cells express NGN3 for less time than WT-NGN3 cells. These differences are illustrated in Fig. 3M.

### Non oscillatory NGN3 results in precocious differentiation to immature endocrine cells and activates target genes earlier

To understand the effect of converting oscillatory NGN3 expression (WT-NGN3) to transiently sustained in PM NGN3 (i.e. one broad peak) we analysed the impact on cell fate decisions. We assessed cell fate by immunofluorescence staining using antibodies against C-PEPTIDE and GLUCAGON at day 23 and day 25 in 3 or 4 independent co-culture differentiations respectively, using mScarleti expression to mark the WT cells (Fig.S2D).

We found that PM-NGN3+ cells show an approximately 2.25x increase in becoming β-like cells (CPEP+/GCG-) and α-like cells (CPEP+/GCG+) compared to WT at day 23 (Fig.4A), which was confirmed by flow cytometry analysis (Fig. S3A,B). This increase seen in differentiation by the PM cells was reduced slightly at day 25 (on average between 1.6 and 2x more than WT cells) (Fig.4A) suggesting an overall shift of the timing of differentiation to an earlier time. The number of delta-like cells (by staining for SOMATOSTATIN) were also investigated using flow cytometry at day 23 (Fig.S3B), but the number of delta-like cells in both WT-NGN3 and PM-NGN3 were very similar to the negative controls (Fig.S3A), therefore it is still unclear whether PM-NGN3 affects the differentiation to delta-like cells, and a protocol that is more focused on generating delta-like cells is needed to answer this question.

**Figure 4.**
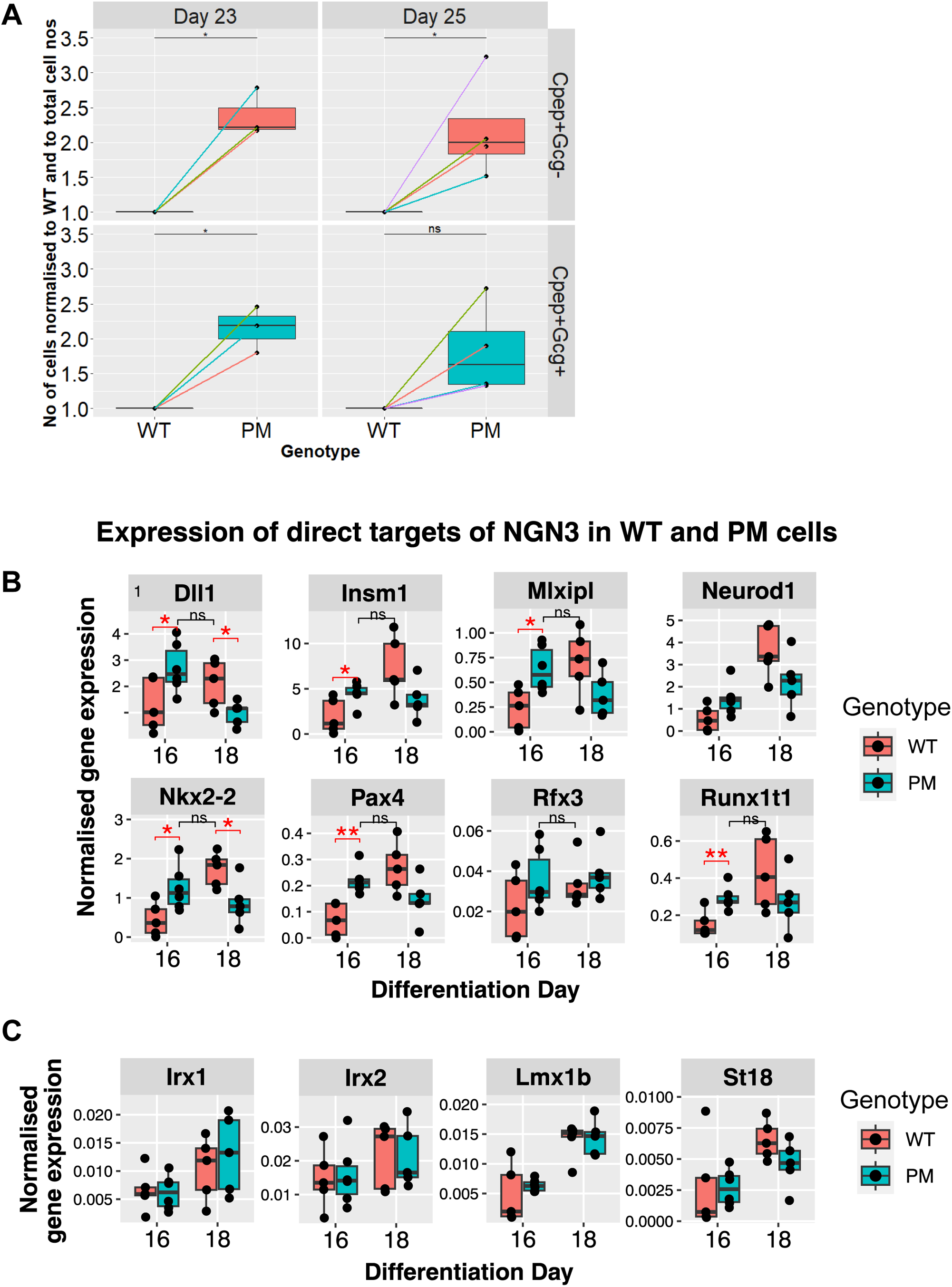
Quantitative analysis of differentiation and NGN3 target gene expression in wild-type and phosphomutant NGN3 expressing progenitors during pancreas differentiation. **(A)** Percentage of wild-type (WT) and phosphomutant (PM) expressing C-peptide (CPEP) with and without Glucagon (GCG) markers compared at differentiation days 23 and 25 from 3 and 4 independent co-culture experiments respectively. **(B) and (C)** Comparative analysis of NGN3 target gene expression levels quantified by RT-qPCR from FACS sorted mVenus positive cells (see Fig. S3C) at differentiation days 16 and 18 in WT versus PM cells; dots indicate independent experiments (5-6), boxes indicate median and interquartile range, statistical significance determined with a t-test for p<0.05* and p<0.01**, p>0.05 non-significant.

To confirm that PM-NGN3 cells are differentiating earlier than WT-NGN3+ cells and to gain some mechanistic insight, we measured the expression of direct NGN3 targets genes at day 16 and day 18 of the differentiation. We performed RT-qPCR from FACS sorted mVenus positive cells (which we show are the cells expressing NGN3 (Fig.S3C)) and we focused on a selection of 12 target genes that: 1) Are bound by NGN3 ^10^, 2) show enrichment in expression in NGN3+ endocrine progenitors ^10^, and 3) have been shown to have important roles in islet cell development ^20–22,60–67^. The resulting shortlist are all transcription factors except for DLL1 which is a NOTCH pathway signaling ligand.

We found that many of these targets (DLL1, INSM1, MLXIPL, NKX2-2, PAX4, RUNX1T1) were significantly increased in their expression in PM-NGN3+ cells at day 16, while most WT-NGN3+ cells did not reach these levels until day 18 (Fig.4B). Indeed, for these genes, there is no significant difference between the expression level between PM-NGN3+ at day 16 and WT-NGN3+ at day 18 (Fig. 4B) suggesting precocious activation of expression. The increases at day 16 in NGN3’s target gene expression in PM-NGN3 cells was found in both separate and co-culture differentiations (Fig. S3D). By contrast, the expression of some other target genes (IRX1, IRX2) was not affected by PM-NGN3 or only showed smaller increases at day 16 in PM-NGN3+ cells (LMX1B, NEUROD1, RFX3, ST18), reflecting that some genes are less able to respond to a change in dynamic expression of NGN3.

In conclusion, non-oscillatory expression of NGN3 results in precocious differentiation to both α-like and β-like cells, most likely by activating important effector NGN3 target genes earlier in the differentiation timeline.

### Time of the maximum fold change of NGN3 correlates with differentiation time

We next wanted to understand how the differences in dynamics of NGN3 in PM-NGN3+ cells can lead to earlier activation of target genes than in WT-NGN3+ cells, and result in differentiation occurring earlier. In this analysis, we used the time of NGN3 turning off as a ‘proxy’ for differentiation, since NGN3 turns off before INSULIN is expressed (Fig. 1F) and PM-NGN3+ turn off NGN3 earlier and differentiate earlier (Fig.3J, Fig.4).

Since NGN3 mainly acts as an activator of transcription ^10^, and given that there is a small (1.3x) increase in maximum NGN3 expression in PM-NGN3+ cells compared to WT-NGN3 cells (Fig. 3H), and PM-NGN3+ cells reach their maximum level earlier than WT-NGN3+ on average (Fig. 3K), one might expect that downstream targets of NGN3 respond to a higher absolute level of expression which drives earlier differentiation. However, we found that neither the median NGN3::mVenus level per cell nor the maximum NGN3::mVenus level per cell, correlated well with the time of NGN3 switching off (differentiation) in both WT and PM conditions (Fig. 5A, S4A, S4C-H). This suggested that the slightly increased absolute level of NGN3 cannot explain why PM cells differentiate earlier.

**Figure 5.**
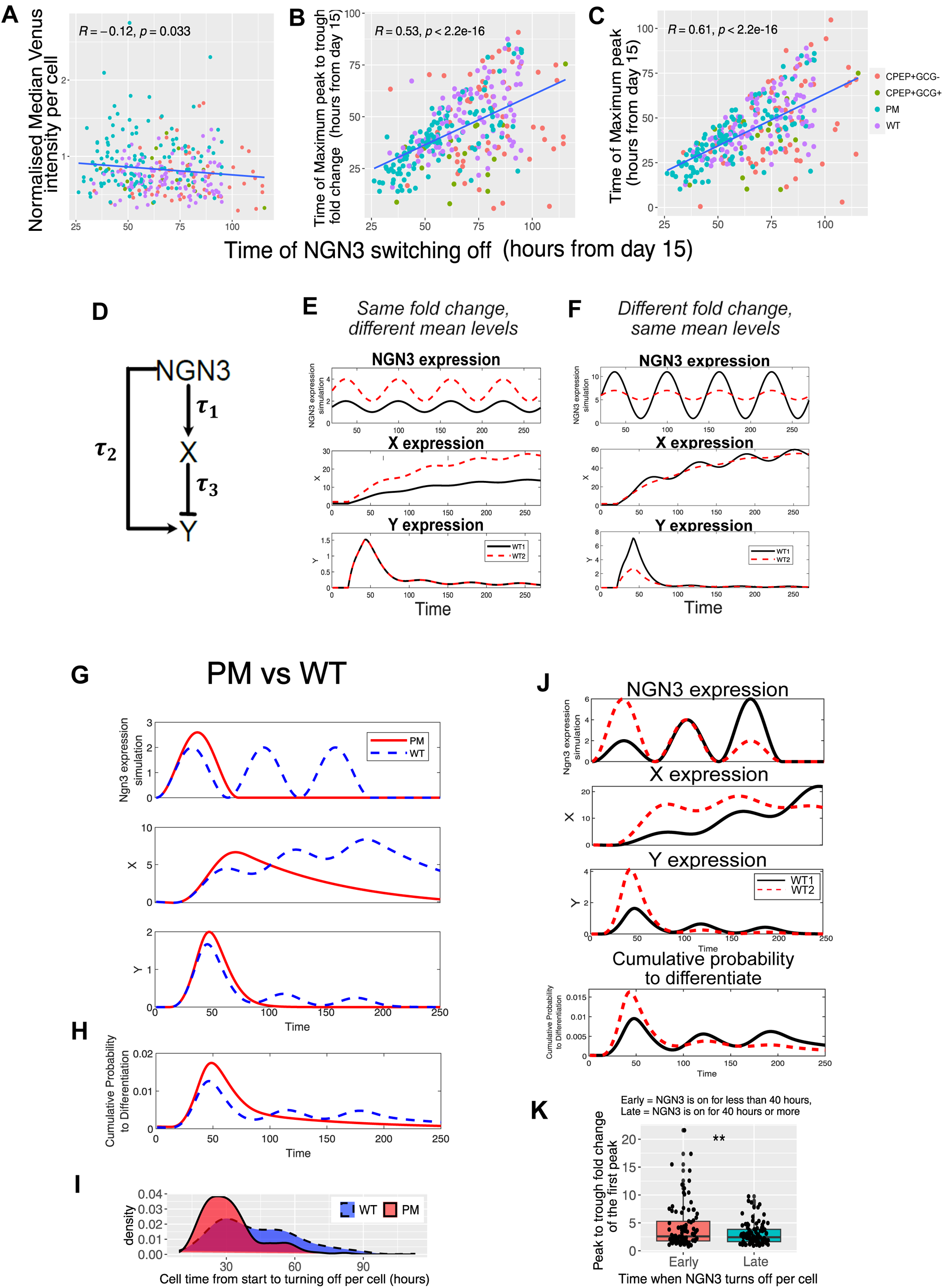
A model of level-independent decoding of NGN3 dynamics by downstream target genes. **(A)** Correlation analysis of normalised median NGN3::mVenus intensity per cell versus the time of switching off NGN3 observed in mixed WT and PM cultures, as well as WT cells that become CPEP+/GCG- and CPEP+/GCG+; in each panel, intensity values were normalised by the maximum of each experiment across 6 independent experiments; R values indicate Pearson’s correlation coefficient. **(B)** Correlation analysis of the time of the maximum peak to trough fold change per cell versus the time of switching off NGN3 observed in mixed WT and PM cultures, as well as WT cells that become CPEP+/GCG- and CPEP+/GCG+; in each panel, intensity values were normalised by the maximum of each experiment across 6 independent experiments; R values indicate Pearson’s correlation coefficient. **(C)** Correlation analysis of the time of the maximum venus intensity value per cell versus the time of switching off NGN3 observed in mixed WT and PM cultures, as well as WT cells that become CPEP+/GCG- and CPEP+/GCG+; in each panel, intensity values were normalised by the maximum of each experiment across 6 independent experiments; R values indicate Pearson’s correlation coefficient. **(D)** The type 1 incoherent feed forward loop (IFFL) regulatory motif encodes how NGN3 oscillations can activate (directly) but also repress a target Y (indirectly -via an intermediate target X). **(E)** Synthetic exploration of motif (D) with target response corresponding to when the input, NGN3 has different mean levels with identical fold-change ratio (2x); X responds to levels of NGN3, in contrast Y is insensitive to a change in level. **(F)** Within the same motif (D), we observe the target response in the case when the input, NGN3 has the same mean level but the fold-change ratio increases from 2x to 5x; X shows a minor difference in response to levels of NGN3 while Y is differentially activated due to fold-change. **(G)** The type 1 IFFL regulatory motif predicts how a difference of 1.15x in fold-change and 1.3x in peak duration observed on average in PM (Red) versus WT (Blue) affects targets X and Y. **(H)** The combination of X and Y responses from (G) to create a probability to differentiate over time in PM and WT simulations. **(I)** Estimation of probability to differentiate from raw NGN3 data observed in PM versus WT illustrated as a density plot using the time each cell expresses NGN3 for before turning off (hours). **(J)** The same motif can be used to explain differences in sinewaves with variable peak heights with one synthetic WT example decreasing from 3x to 2x to 1x (Red-WT1) and another increasing from 1x to 2x to 3x (Black-WT2); the response in X and Y are affected with the larger fold-change at the start of WT1 causing both A and Y to increase dramatically, whereas in WT2 X accumulates gradually and Y responds to each pulse of NGN3. The combined probability to differentiation due to X and Y is shown in the bottom panel. **(K)** Comparative analysis of initial peak to trough fold-change values in WT cells that downregulate NGN3 early (NGN3 is on for less than 40 hours) versus late (NGN3 is on for more than 40 hours). Statistical significance determined with a t-test, p<0.01.

There was also poor correlation between the maximum peak to trough fold change per cell and the time of NGN3 turning off (differentiation) (Fig.S4A), however, we found that there was good positive correlation between the time that the cell reached this maximum fold change and the time of differentiation (Fig. 5B, S4A). There was also a similarly good correlation between the time the cell reached its maximum level (Fig.5C, S4A) and differentiation, reflecting the good correlation between maximal fold change and maximal level (Fig.S4A,B). Taken together, these observations suggest that NGN3 may affect the timing of differentiation (quantified here as turning off of NGN3) through the timing of a change in fold change, i.e. a maximum peak to trough, rather than absolute median or maximal level of NGN3 expression.

### An incoherent feed forward loop (IFFL) explains the premature differentiation of altered, non-oscillatory, NGN3+ dynamics

To understand how a mechanism that detects changes in fold-change would work we turned to theoretical work on an IFFL ^68–70^ a powerful and widespread GRN motif which can allow the fold change of the input to be decoded rather than its absolute levels. The principle of an IFFL is that an input activates a target as well as repress it ^68,70^. In the type 1 incoherent feed-forward loop (I1-FFL) regulatory motif used here, an upstream gene (here, NGN3) can activate its target Y but also repress it through the activation of an intermediate repressor, X (Fig. 5D). A fold change activated target can increase its level in response to oscillatory expression, a single pulse, or a step increase of the input. Thus, to understand fold-change detection in the context of this work, we studied the I1-FFL regulation firstly under imposed oscillatory activation (a similar idea has also been studied in ^71^). In this scenario, a level dependent target, such as X, ‘integrates’ the history of NGN3 activation leading to accumulation of X over time (if the protein is sufficiently stable (Fig. 5E, S4I)), and different mean levels of the oscillatory NGN3 input result in more X produced over time (level-dependent decoding). At the same time, the fold-change dependent target, Y, transiently increases until the level of X represses Y causing it to downregulate, giving rise to a pulse of Y (Fig. 5E,F). Different NGN3 inputs with the same fold-change will produce identical responses from Y irrespective of NGN3’s absolute levels (Fig.5E), i.e. the same response from Y will be produced from inputs of, for example, a signal at 10 A.U. which jumps to 20 A.U. as it will for one at 500 A.U. jumping to 1000 A.U. In comparison, if we use inputs with the same mean levels but different fold-change, then this results in a very similar response in X, but the increase in fold change increases the production of Y directly before it can be repressed by X, resulting in a peak in Y that decodes different fold-changes (Fig. 5F). Other target dynamics are also possible depending on the stability of X and Y (Fig. S4I), however in all examples the fold-change dependent target Y is insensitive to the level of NGN3 (Fig. 5E, S4I). Thus, while both X and Y are directly induced in a level dependent way by the input (NGN3) and the amount of X produced is NGN3 level-dependent, the amount of Y produced is NGN3 level-independent but is responsive to the fold change of NGN3’s oscillations.

To understand how the time of the largest fold-change correlates with differentiation from our data (Fig 5B, S4A) we interrogated this motif comparing WT-NGN3 oscillatory activation with PM-NGN3 pulse activation imposing inputs that resembles the data (Fig. 5G). We observed that Y would be activated before X in both WT and PM and that the height and duration of the Y response is higher and longer in PM-NGN3 cells likely due to PM-NGN3’s increased fold change and peak duration. At the same time, activation of X would initially be higher in PM-NGN3 cells, however WT could achieve the same level of X later by incrementally accumulating with every subsequent peak of WT-NGN3. If we assume that activation of both X and Y targets increases the probability to differentiate, i.e. that differentiation is achieved by the combinatorial activation (Fig. 5H), differentiation of PM-NGN3 cells would occur earlier, while in WT-NGN3 cells, oscillatory NGN3 would moderate and spread out this probability over time. This agreed well with the experimental data where the distribution of the time of switching off occurs earlier and more synchronously in PM-NGN3+ compared to WT-NGN3+ cells (Fig. 5I).

### An IFFL also explains the spread in differentiation of wild-type oscillatory NGN3+ dynamics

To achieve a unifying mechanistic understanding, we asked whether the model above would also be able to explain the variation in the time of differentiation among WT-NGN3 expressing cells (Fig 3J). In WT-NGN3+ cells, there are multiple NGN3 peaks, with varying fold change between peaks (Fig. 1G, 1H, 2A). In the model, by varying NGN3’s fold change over time, the expression of Y and X targets are affected, changing the probability of differentiation across time (Fig. 5J). A large fold change at the beginning of the cell’s NGN3 expression window, allows for both X and Y to reach high levels quickly, creating a large probability to differentiate at an early time point (red dashed line in Fig. 5J, similar to the PM-NGN3+ cell simulation). In comparison, a cell with a small fold change at the beginning which increases with every peak over the NGN3 expression window, results in a slow accumulation of X and multiple smaller peaks of Y (due to less X produced, and therefore less repression of X onto Y)(Fig. 5J). This results in the cell being equally likely to differentiate early or late (Fig. 5J). We hypothesize that if the cell was unable to activate enough Y through a fold-change mechanism during the first peak of NGN3, multiple peaks of NGN3 or subsequent large fold change peaks will activate level-dependent X targets to increase the probability to differentiate (black line in Fig. 5J). Indeed, in the experimental data, we have found that WT cells which turn off NGN3 earlier tend to have a first peak with a larger fold change than WT cells that turn off later (Fig. 5K), supporting the existence of this mechanism in WT cells. Such a mechanism could provide an advantage to WT cells enabling cells to spread out their differentiation over time without prematurely depleting the progenitor pool, and creates robustness in the system by allowing differentiation to occur through the balance of multiple target genes, and the decoding of both levels and fold-change in the input.

In conclusion, by proposing a theoretical I1-FFL fold change-based decoding, we were able to explain the distribution of NGN3 switching off timings in cells with oscillatory NGN3 expression (WT) as well as the premature differentiation observed in cells with a single broad peak of NGN3 expression (PM). In addition, our synthetic exploration predicts the existence of targets downstream of NGN3 with different dynamics, of either continuous activation (resembling X) or a peak of expression (resembling Y) over time, which we examined next.

### Bioinformatic analysis supports the existence of a NGN3 controlled IFFL

To examine if this gene network motif exists in human pancreatic development, we first investigated whether we saw differences in expression dynamics of the direct targets in this differentiation using RNA-seq. Focusing on the genes bound by NGN3 ^10^ and upregulated once NGN3 is expressed on day 15 compared to day 12, we found the presence of both accumulating (cluster 1) and transiently expressed (cluster 2) targets corresponding to X and Y profiles of the model respectively (Fig. 6A, B, C). As our theoretical model shows, the I1-FFL motif can explain the emergence of these two dynamics from the same upstream oscillatory activation (NGN3).

**Figure 6.**
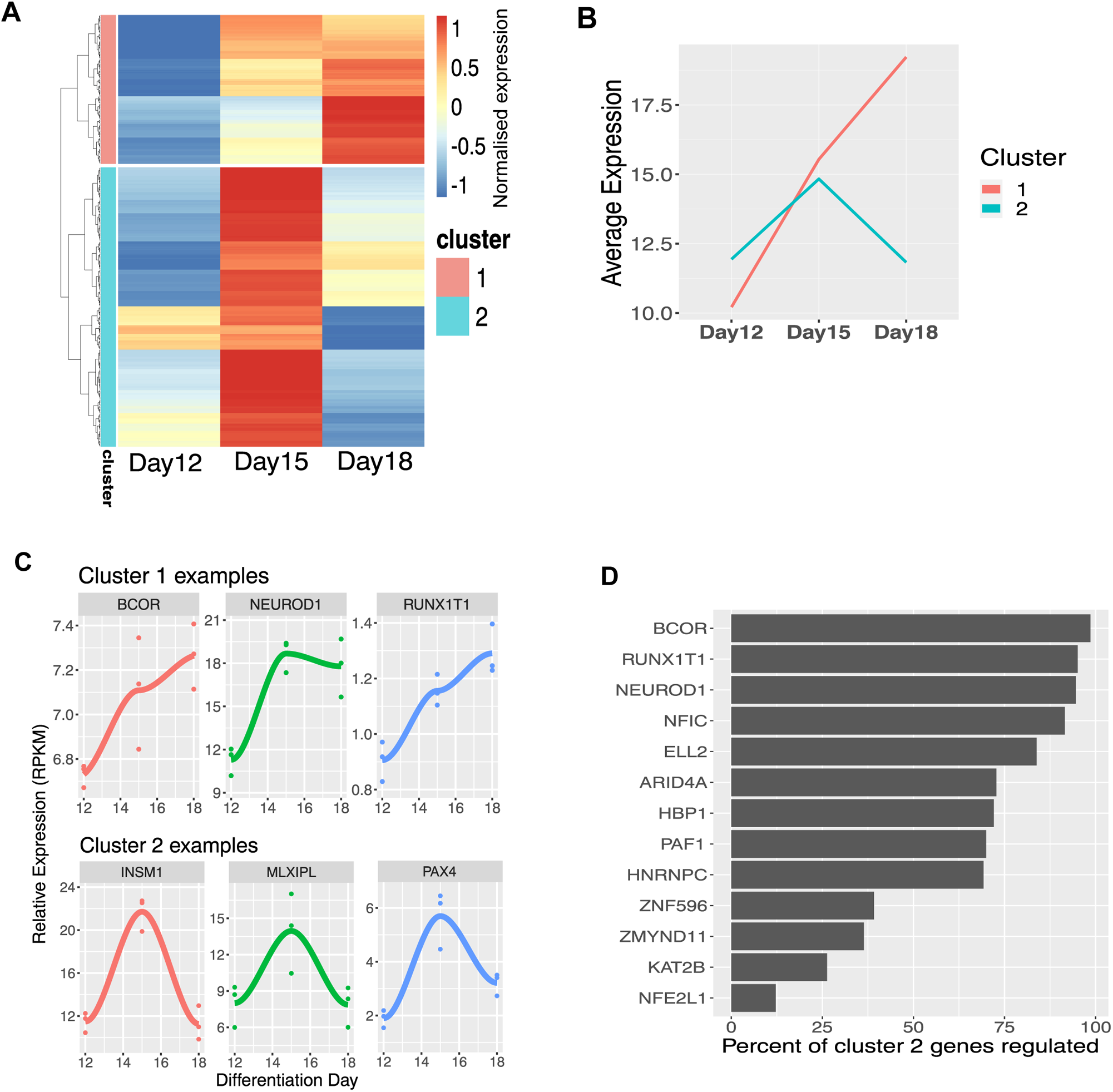
Bioinformatic exploration of NGN3 target gene dynamics and interactions. **(A)** Clustering of z-scored RPKM target gene expression values at differentiation days 12, 15 and 18 quantified using bulk RNA sequencing reveals two main cluster dynamics. Rows are genes (see methods) and columns are differentiation day time points samples were taken. **(B)** Average expression per cluster using RPKM values from the RNA-seq data indicates two distinct profiles with cluster 1 increasing over time and cluster 2 showing a peak of expression. **(C)** Gene expression profiles for individual genes corresponding to cluster 1 (top panel) and cluster 2 (bottom panel). **(D)** Percent of cluster 2 genes bound by each of the 13 cluster 1 genes present in the ReMap ChIP-seq dataset (Materials and methods).

To further confirm that it is likely these targets genes could be part of this motif, we used the ChIP-seq data from ReMap ^72^ to look for potential interactions between genes in cluster 1 and cluster 2, as the fold-change detection relies on the repression of targets Y by targets X. Of the cluster 1 genes, 13 of these were present in the ReMap data, and they all show binding sites associated with cluster 2 genes; ranging from binding 52 of cluster 2 genes (12.2% of all cluster 2 genes) to binding 420 of cluster 2 genes (98.6% of cluster 2 gene) (Fig. 6D). While ChIP-seq binding data does not indicate whether these interactions are activatory or repressive, BCOR (binds 98.6% of cluster 2 genes) and RUNX1T1 (binds 95.1% of cluster 2 genes) are both known repressors ^73–75^. NEUROD1 (binds 94.6% of cluster 2 genes) however, is more complicated since it has been shown to act as an activator as well as repressor ^76^, but it is possible that the I1-FFL motif is formed by the combined interactions of the target genes.

## Discussion

In this paper, we have studied the dynamics of NGN3 expression, a transcription factor important for pancreatic development in an iPSC-based model of human pancreatic development. We reasoned that the precise protein dynamics, as revealed with endogenous reporters, at the single-cell level and with live imaging, encode information which is not accessible with either snapshot or population based measurements. We have asked what are the precise dynamics of the endogenous human NGN3 protein expression, whether the dynamics are functional and how are they decoded.

We report that the protein expression of NGN3 is oscillatory in human pancreatic endocrine progenitors within a time window of a few days (in vitro) and is switched off upon differentiation. Oscillations of the TF HES1 and the signalling molecule DLL1 have recently been reported in murine pancreatic development and these oscillations are important for proliferation in the non-endocrine lineages ^33^. Our finding that NGN3 oscillates represents the first oscillatory TF to be found in human pancreatic development and suggests that sub-circadian oscillations (i.e. ultradian) play a role in human development. The regulatory region of the related helix-loop-helix gene *Ngn2* has been shown to drive Ub-Luc oscillations in mouse cortical progenitors, denoting broad conservation of mechanism both across species and across tissue organs. However, there are also salient differences; murine *Ngn2* oscillates in cortical neural progenitors and is sustained at a high level in differentiating cells, while human NGN3 oscillates in post-mitotic pancreatic endocrine progenitors and is switched off upon differentiation. In addition, the periodicity of the human NGN3 protein reported here (13hr) is longer than the one reported for the mouse *Ngn2* (4-6hrs; Fig. 7D in Shimojo et al.,2008 ^77^), most likely reflecting the slower tempo of human development^78^, as it has been shown to be the case for somitogenesis associated oscillators^79,80^.

What is the function of NGN oscillations? In mouse neurogenesis, the transition of oscillatory to sustained expression of distinct transcription factors has been associated with the choice of different fates giving rise to the notion that oscillations may be decoded in the choice of different fates ^27^. Consistent with this, during spinal cord differentiation different HES1 dynamics (oscillatory compared to aperiodic) correlated with a fate choice^24^. However, in the case we describe here, the characteristics of NGN3 oscillatory expression (fold change, period, mean level and duration of oscillations) are the same in progenitors that lead to the two main pancreatic cell fates produced by this system, β-like and α-like cells, suggesting that oscillations are not decoded in differential fate acquisition. This is in agreement with previous molecular studies ^10^ which showed that NGN3 does not directly bind specific hormone genes but instead regulates pan-endocrine genes, therefore unlikely to directly influence cell fate. In addition, Mastracci et al., 2013 ^81^ suggests that fate specification may occur before NGN3 expression. Therefore, we hypothesize that NGN3 oscillatory expression induces differentiation but does not affect the specific cell fate choice. It is formally possible though that NGN3 dynamic expression plays a role in the specification of other hormone producing cells (e.g. SOMATOSTATIN expressing cells) which could not be assessed with the current differentiation protocol.

To gain insights into the functional role of oscillatory NGN3 expression, we experimentally manipulated NGN3 expression dynamics by mutating the C-terminal phosphorylation sites. This modification (PM-NGN3) increased the stability of NGN3 protein, as it has been previously described both for mouse and human NGN3^36,58^, and resulted in the majority of cells expressing NGN3 as one longer peak of expression in place of multiple shorter duration peaks (oscillations). Under these conditions, we observed accelerated differentiation to both β-like and α-like cells, preceded by premature activation of important target TF genes (e.g. *NKX2-2*, *PAX4*, *MLXIPL*, *RUNX1T1*), without biasing towards either fate. Previous work showed an increase in α- and δ-cells in mice expressing PM-NGN3, assessed as a snapshot during the secondary transition of differentiation^36^. To reconcile these findings and based on the time resolution afforded by our studies, we suggest that the apparent increase in differentiation reported in Azzarelli et al., 2017^36^ may represent differentiation that has occurred earlier in the PM-NGN3 expressing mice. Knowledge of NGN3’s natural oscillatory dynamics, and the resultant change of dynamics in the PM version, complements a previous report that the PM murine NGN3 has a greater affinity to bind DNA^36^.

What is the mechanism by which NGN3 dynamics impact the time of differentiation? To answer this question, we sought to explain both the premature differentiation elicited by the one long peak of PM-NGN3 expression as well as the natural temporal spread of differentiation observed in cells where WT-NGN3 oscillates. Motivated by the wide range in the level of NGN3 expression in WT cells and the increase of expression, albeit modest, (1.3x) in PM cells, we examined the influence of the level of expression. However, we have found no correlation between the mean and maximal level of expression with the timing of differentiation in either PM or WT NGN3 expressing cells, suggesting that the absolute NGN3 level is not a good predictor for the timing of differentiation. Instead, we did find a good correlation between the time of the maximum fold change, measured between trough and peak, and the timing of differentiation. This finding suggested that a large fold change observed in single-cell expression timeseries of either the WT or the PM NGN3, can initiate the gene expression changes that are needed for differentiation to proceed, and the timing of this large fold change is instructive for differentiation.

In order to gain insights into how the timing of a fold change difference may be decoded, we focused on the possible involvement of an IFFL motif downstream of NGN3. We explored a particular type of such motif where a downstream target would be activated by NGN3 but also repressed (with delay) due to activation of an intermediate repressor. When IFFLs are employed for signal decoding, the output shows a transient peak that returns to the pre-stimulation level, a process known as perfect adaptation and important for homeostasis^69^. Fold-change detection through an IFFL have also been described as a robust method of signal decoding since it buffers against stochastic variation in absolute gene expression levels that may naturally occur from cell to cell. In addition, signal decoding via an IFFL would give a transient output peak before it returns to basal level, but this peak would have the same characteristics if the fold change of the input is the same and irrespective of its level. Indeed, there is evidence that fold changes are read in the ERK, NODAL, NFκB and WNT signalling systems ^45,82–84^. This is particularly important for NGN3 where cells show a widespread variation in the level of expression, yet there is no correlation with the timing of differentiation. While most studies have focused on the ability of the IFFL to interpret and give the same output at different levels but same fold change of the input, here, we have additionally explored the involvement of an IFFL when the fold change of the input differs. Indeed, our experimental data show that the fold change of WT-NGN3 expression differs over time and that PM-NGN3 consistently shows a higher fold change. In both genotypes, we see a positive correlation of the timing of the higher fold change with the timing of differentiation, consistent with an altered response of the IFFL system to a fold change difference. Thus, a widespread (and underappreciated) mechanism of decoding oscillations is through fold change detection.

Taken together our data shows that NGN3 has an additional dimension in its expression in human pancreatic endocrine progenitors in that it shows oscillations with ultradian periodicity. During human development the window for pancreatic endocrine differentiation via NGN3 spans across at least a 3-month period^39^ highlighting the likelihood for mechanisms to be in place to be able to differentiate throughout this time window. Our work suggests that that NGN3 oscillations are normally exploited to spread out the timing of differentiation in a population of pancreatic endocrine progenitors, therefore differentiation events are temporally asynchronous. This might also suggest why in vitro differentiation protocols are not very efficient^85–87^, since in these protocols the expression window is only 4-5 days long, and perhaps not long enough for all oscillatory NGN3 expressing cells to differentiate. These findings are important for more precise engineering of pancreatic differentiation.

## Author Contributions

A.M and N.P conceived and designed the experimental study. A.M generated cell lines, performed pancreatic endocrine differentiations, imaging, immunofluorescence and flow cytometry analysis, acquired, tracked and analysed single-cell imaging movies, analysed RNA-seq data, performed RT-qPCR, wrote the paper. V.B analysed dynamic expression from single-cell imaging movies (percent oscillators, period, peak to trough fold change, percent of cells with one peak) co-supervised A.K with A.M, and collaborated alongside A.M with A.R on the fold-change motif, and wrote methodology for the paper. A.R implemented a fold-change detector motif model and and wrote methodology for the paper. F.W performed half-life experiments and analysed the data under supervision of A.M and wrote methodology for the paper. M.C performed the differentiation experiments that were sequenced for the RNA-seq data under the supervision of L.V. A.K created time-series data from imaged single-cell time-lapse movies under the supervision of A.M and V.B. E.M performed immunofluorescence on fixed differentiation samples. N.P supervised the work, interpreted data and co-wrote the paper with A.M with input from V.B.

## Acknowledgements

The authors would like to thank the Bioimaging facility (especially Dr. Peter March and Dr. David Spiller) and the Flow Cytometry facility (especially Dr. Gareth Howell) of the University of Manchester, Dr. Cerys Manning for reading and providing comments on the manuscript and all Papalopulu lab members for comments and discussions. The work was supported by a Sir Henry Wellcome Trust Postdoctoral Fellowship to A.M (210912/Z/18/Z) and a Wellcome Trust Senior Research Fellowship to N.P (224394/Z/21/Z).

## Declaration of interests

The authors declare no competing interests.

## Methods

### Maintenance of hIPSc

FSPS13b hIPSCs (http://www.hipsci.org/lines/#/lines/HPSI0813i-fpdm_2, RRID:CVCL_AH05) were cultured in Essential 8 or Essential 8 FLEX media (Life Technologies # A1517001 and # A2858501) on vitronection (Thermo Fisher #15134499) coated plates. All tissue culture was carried out in 5% CO_2_ at 37°C. The medium was changed every day in Essential 8, or every two days in Essential 8 FLEX, and cells were passaged every 4-5 days using 0.5 mM EDTA (Life Technologies, #15575-020) to dissociate cells maintaining clumps. In all hiPSC cultures, 10 µM Rho-associated protein kinase (ROCK) inhibitor, Y-27632 (Bio-techne #1254/10), was only added into the culture media when thawing hiPSCs. Human iPSCs were maintained at 37°C with 5% CO2 and regularly tested negative for mycoplasma contamination.

### Generation of WT-NGN3::mVenus and PM-NGN3::mVenus clonal hIPS lines

CRISPR/Cas9 RNP protocol was followed as described in ^88^. The HDR template was designed as follows: Linker-3xFLAG-mVenus-HA with 800bp either side of homology arms to NEUROG3 gene. Guide RNA targeting the NEUROG3 locus (TTTTCCGCCTGCTTGAGCCC-AGG) was purchased as crRNA (IDT) and annealed with tracrRNA (IDT) in duplex buffer provided at 95°C for 5 minutes then cooled slowly to room temperature before mixing with Alt-R™ S.p. HiFi Cas9 Nuclease V3 protein (IDT) and incubated for 20minutes at room temperature. 800,000 cells were resuspended in 100ul of Lonza P3 Primary Cell Nucleofector buffer, and HDR template (2µg) and Cas9:gRNA mix (final concentration of 2.5µM) was added, and electroporated using pulse code CA-137. Cells were plated onto Synthemax coated plates (5 μg/cm2) in Essential 8 with 10 μM ROCK inhibitor, and media was changed to Essential 8 24 hours later. Cells were plated for single cell clones by plating 1000 cells per 10cm^2^ dish on Synthemax with 1x CloneR (Stemcell technologies #05888), and single cell clones were genotyped by PCR for correct integration before sequencing both alleles from positive clones.

To create the PM-NGN3::mVenus cell line, site directed mutagenesis was performed on the NEUROG3::mVenus HDR plasmid template to mutate the 5 alanine residues (Fig. 3A) using NEB Q5 site directed mutagenesis kit (#E0554S) following manufacturer’s instructions. mVenus and the mutations were integrated into the NEUROG3 locus using CRISPR/Cas9 HDR as above. Since all 5 mutations were very close to the double strand break site (all within 132bp upstream) if HDR occurred and mVenus was integrated, it was also likely the mutations were “repaired” as well. Genotyping of single cell clones was performed by first checking for the integration of mVenus by PCR and then both alleles were sent for sequencing to validate for the 5 mutations. Of the 11 single cell clones that had the correct integration of mVenus, 4 had all the 5 mutations also integrated (36% success rate), while another 4 had only repaired the closest mutation to the cut site, and the remaining 3 clones has mVenus integrated but none of the mutations. Only one cell clone was homozygous for mVenus and all 5 mutations, so this clonal cell line was used for all subsequent experiments.

### Generation of NGN3::mVenus + NLS-mScarleti-NLS clonal line

Homozygous WT-NGN3::mVenus (800k cells) was nucleofected using Lonza P3 primary cell kit (pulse code CA-127) and 2µg of CAG-NLS-mScarleti-NLS-IRES-Puromycin plasmid, plated onto 2x Vitronectin coated 6 well plates in Essential 8 media plus 10µM ROCK inhibitor. 24 hours later, media was refreshed with Essential 8 and 0.2µg/ml puromycin. Puromycin was maintained in the media for at least 5 passages. Once cells were selected, single cell cloning was performed to create a line with similar expression levels of the Scarlet transgene. The cells were single cell sorted by FACS based on Scarlet expression into a Synthemax coated 96 well plate (5 μg/cm2) in Essential 8 plus CloneR, and one line was carried forward for experiments.

### Differentiation to pancreatic endocrine

Pancreatic differentiation was carried out as in El-Khairi et al., 2021^47^, as follows (N.B all incubations were carried out at 37°C 5% CO2). Tissue culture dishes were coated by incubating overnight at with 0.2% gelatin in PBS, followed by the removal and replacement with Advanced DMEM F12 + 10% FBS overnight in the incubator. The following day, this was removed and replaced by single hIPS cells in Essential 8 or Essential 8 Flex plus 10µM ROCK inhibitor. Cells were plated at 22,000cells/cm2. The following day, media was refreshed with Essential 8 or Essential 8 FLEX only (this is day 0). On day 1, cells were cultured in CDM-PVA or CDM-BSA supplemented with Activin A (100ng/ml), FGF2 (80ng/ml), BMP4 (10ng/ml), CHIR99021 (3μM) and Ly294002 (10μM) and Day 2 with CDM-PVA or CDM-BSA supplemented with Activin A (100ng/ml), FGF2(80ng/ml), BMP4 (10ng/ml) and Ly294002 (10μM). On Day 3 RPMI/B27 media containing Activin A (100ng/ml), FGF2 (80ng/ml). Days 4-6, the cells were cultured in Adv-BSA media supplemented with SB-431542 (10μM), FGF10 (50ng/ml), Retinoic Acid (RA; 3μM), Noggin (150μg/ml) and L-Ascorbic acid (0.25mM) with freshly made media changed daily. For days 7-8, cells were cultured in Adv-BSA with FGF10 (50ng/ml), RA (1μM), Noggin (150μg/ml), KAAD-cyclopamine (0.238μM), PdBU (50nM) and L-Ascorbic acid (0.25mM) with media changed daily. Days 9-13 cells were cultured in RA (100nM), Noggin (150μg/ml), KAAD-cyclopamine (0.238µM), EGF (100ng/ml), Nicotinamide (10mM), and L-Ascorbic acid (0.25mM) with media changed every 48hours. On Day 14 media was replaced to Adv-BSA plus glucose (final concentration 25mM), B27 (1%), RA (100nM), DAPT (1μM), Alk5i (10μM) and 6-Benzoyladenosine-3’,5’-cyclic monophosphate (BNZ,0.1mM) for 3 days. On Day 17 media was replaced with Adv-BSA containing B27 (1%), RA (100nM) and Alk5i (10μM), and from day 20 onwards media was replaced with Adv-BSA containing B27 (1%), RA (100nM) and refreshed every 3 days until required.

Basal medias mentioned above: CDM-PVA (1:1 mix of F12 and IMDM plus 0.1% PVA, 1% Conc lipids, 0.004% 1-Thioglycerol, 7ug/ml Insulin, 15ug/ml Transferrin, 1x Pen/Strep), CDM-BSA (1:1 mix of F12 and IMDM plus 0.5% BSA, 1% Conc lipids, 0.004% 1-Thioglycerol, 7ug/ml Insulin, 15ug/ml Transferrin, 1x Pen/Strep), RPMI/B27 (RPMI 1640 + GlutaMax plus 2% B27, 1x NEAA, 1x Pen/strep, Adv-BSA (Advanced DMEM/F12 plus 0.5% BSA, 1x Pen/Strep, 1x GlutaMax).

### Time-lapse imaging and single cell tracking

Time-lapse movies were generated by plated the differentiations in 35Lmm glass-bottomed dish (Greiner BioOne), ibiTreat u-dish 35mm High (Ibidi) or ibiTreat u-dish 35mm high Grid-500 (ibidi). The addition of HEPES (Life tech #) to a final concentration of 25mM, was added into any media that was added onto the cells while in the microscope. Movies were acquired using Zeiss LSM880 microscope and GaAsP detectors. A Plan-Apochromat ×20 0.8 NA objective or Fluar ×40 1.3 NA objective were used. Z-sections covering the depth of the cells were acquired every 10 or 15 min for 48-144Lh. Max projections of timelapse movies were created with Fiji (RRID:SCR_002285), and then single cell tracks were produced in Imaris (RRID:SCR_007370) using the ‘Spots’ and ‘Track over time’ function. All tracks were manually curated to ensure accurate single-cell tracking, and generated mean intensity over time, representing protein concentration over time in single cells.

### Immunofluorescence

Cells were fixed using 4% PFA for 15 minutes followed by a PBS wash. The cells were then permeabilised and blocked using PBS + 0.1% triton X + 10% donkey serum for 1 hour at room temperature. Primary antibodies (Table S1) were diluted in PBS + 0.1% triton X + 1% donkey serum and incubated on the samples overnight at 4°C. Three 10 minutes washes in PBS was then performed and then secondary antibodies (Table S1) added, diluted in PBS + 0.1% triton X + 1% donkey serum, incubated at room temperature for 1 hours, kept in the dark. Three more PBS washes were then performed, and left in PBS for imaging.

### Fate Mapping

To monitor the NGN3 dynamics of single cells and then subsequent fate, the WT-NGN3::mVenus cells with NLS-Scarleti-NLS expressed were used to be able to keep track of the cells beyond the NGN3 expression window. These were differentiated and imaged as described above, on ibiTreat u-dish 35mm high Grid-500 (ibidi) dishes, and at day 19 or day 20 of the differentiation, removed from the microscope and fixed and stained immediately (as described above).

### FACS and qRT-PCR

Cells were sorted on a BD Influx and cells positive for mVENUS expression were sorted directly in 1ml TRIzol reagent (Thermo Fisher Scientific, Cat# 15596018) that was kept on ice. 200μl of chloroform were then added to each sample and samples were centrifuged at 14,000rpm for 15 min at 4°C. The clear phase of the solution was transferred into a new Eppendorf tube to which 500μl of isopropanol and 1ul of GlycoBlue (15mg/ml) (Thermo Fisher Scientific, Cat# AM9515) were added. Samples were kept at −20°C overnight and then centrifuged at 14,000 rpm for 15 min at 4°C. The supernatant was removed and pellets were washed with 70% ethanol and then centrifuged at 14,000 rpm for 10 min. Pellets were reconstituted in RNAse-free water and treated with DNase-I (Promega ##) as per manufacturers’ instructions. cDNA was prepared using Superscript III (Invitrogen, Cat# 18080051) as per manufacturers’ instructions and qPCR was performed with SYBR green (Applied Biosystems, Cat# 4367659) or Taqman (Cat #4453320) using the primers and probes listed in Table S2, Table S3. CT values were normalized to reference genes ACTB or RPL7.

### Flow cytometry

Cells were removed from the cell culture plates and created a single cell suspension using accutase. Cells were fixed in 4% PFA for 15 minutes, spun at 300g for 3minutes to pellet the cells and then resuspended to permeabilised with PBS + 1% Saponin for 30 minutes at room temperature. Cells were centrifuged again, supernatant removed, then resuspended in PBS+ 5% FBS + 1% Saponin plus primary antibodies and incubated for 1 hour at room temperature. Two washes in PBS + 5% FBS + 1% saponin were then performed, and the secondaries were diluted in PBS + 5% FBS + 1% saponin and incubated for 30 minutes at room temperature. Two more washes in PBS + 5% FBS + 1% Saponin, then cells were left in PBS + 2% FBS for flow cytometric analysis. All samples were run on the BD LSRFortessa™ Cell Analyzer and the data were analysed with FlowJo 10.8.0 (RRID:SCR_008520)

### Half-life experiments

FSPS13b hiPS cells were nucleofected using the P3 Primary Cell Nucleofector^TM^ Solution (Lonza #V4XP-3024) with pulse code CA137. 800k cells were nucleofected with 2ug of a plasmid constitutively expressing either Ngn3-linker-3xFLAG-Dendra2-HA, PM-NGN3-linker-3xFLAG-Dendra2-HA or Dendra2 alone. The cells were plated onto glass bottom 35mm dishes (Greiner #627871) in Essential 8 FLEX media and 10uM ROCK inhibitor and incubated at 37°C and 5% CO2 for 24 hours before imaging. Time-lapse movies were acquired using Zeiss LSM880 microscope and GaAsP detectors using a Fluar ×40 1.3 NA objective. The Dendra2 fluorescent protein was photoconverted using UV light for 5 seconds, and then the photoconverted version of the protein was imaged every 2.4-4.35 minutes. Max projections of time-lapse movies were created with Fiji (RRID:SCR_002285), and then single cell tracks were produced in Imaris (RRID:SCR_007370) using the ‘Spots’ and ‘Track over time’ function, creating fluorescent intensities over time of the photoconverted red Dendra. To calculate the protein half-lives from the temporal decay of red Dendra emission we plotted exponential regression lines. Background fluorescence was removed from cell traces, and the plateau of the regression was set to 0. To account for photobleaching cell traces were normalised to the Dendra2 alone control. When the normalised fluorescence intensities per cell over time were plotted, the extra sum-of-squares F test showed an individual exponential regression curve per genotype had a significantly better fit (P<0.0001) than a single curve for all experiments. We therefore calculated the half-lives per genotype per experiment using the one-phase decay equation in Prism. To test if the half-lives across all experimental repeats were significantly different from each other, the Shapiro-Wilk normality test showed our data was normally distributed, and therefore, the paired t-test was used.

### Detection of Oscillations by a Gaussian Process with local detrending

To detect oscillatory activity from single cell timeseries (or tracks) we used customisable Gaussian Processes pipeline previously described in ^24,56^. These approaches were designed for short timeseries spanning 2 to 3 peaks, however in this study individual cells were tracked for several days leading to overall longer tracks and inherent variability in track length (spanning 22h to 55h, see Table S4). Long-term imaging introduced a transient artifact around changing of media which could be wrongly interpreted as a trough in individual cell intensity; to prevent this we split the tracks in before and after the time of media change and we analysed the resulting partial tracks individually. The first step in analysing the oscillatory activity is detrending, a procedure that improves the local estimation of period duration and periodicity by removing variability in mean level occurring at longer timescale than 2 to 3 times the period of the oscillation. Here we used a 20h time-window to fit linear slow variations in mean level, referred to as trend and subtracted this to produce detrended timeseries (see examples in Fig S1H). Following detrending, the timeseries were analysed using a Gaussian Process with a wave-like covariance model and its likelihood was compared to the likelihood of a covariance model with aperiodic fluctuations by means of the log-likelihood ratio statistic (LLR, see^56^). The distribution of LLR from all timeseries in a single experiment was compared to synthetic aperiodic data via a 5% false discovery rate test, thus assigning an oscillatory or non-oscillatory label to each track. In this improved detection method, we assessed that track length does not affect the LLR and period estimation (see Table S4, correlation tests).

### One peak auto-correlation testing with statistical significance

The ability of Gaussian Process models to discriminate between persistent oscillatory activity and a single peak dynamic is poor since these dynamic types would always outperform the aperiodic fluctuating covariance model. To overcome this limitation, we used auto-correlation function (ACF) analysis which measures similarity of a signal to the same signal shifted by different time duration (relative or lag time). In persistent oscillatory timeseries, the autocorrelation function has multiple peaks indicative of multiple oscillatory waves re-occurring over time; however, in the one peak dynamic type resembling PM, the autocorrelation function has a single peak at lag 0, followed by a dip and a plateau. To determine the percentage of timeseries with one peak, we identified ACF peaks and plateaus above 1.5 standard deviation of noise, corresponding to a 90% significance level. Non-significant peaks at less than 1.5 SD were rejected thus ensuring non-oscillatory timeseries are excluded from the analysis. To avoid spurious results due to short track length, we excluded experiments with less than 30h average length from this analysis. The ACF approach enabled us to measure individual peak durations regardless of dynamic type. We report peak duration as the time interval required for the ACF curve to reach the first peak or plateau.

### Peak to trough fold change

We quantified the peak to trough fold changes of the NGN3::mVenus signal from raw undetrended timeseries. Unlike amplitude which is typically measured from the mean level and expressed in arbitrary intensity units, the fold change ratio can be compared across experiments as it is unitless. With the completion of every oscillatory wave, the instantaneous phase angle (obtained from the Hilbert transform) resets ranging values between -π and + π. The peaks and troughs in the signal correspond to points where the phase angle crosses the value 0 and we exploited this property to detect them. Every detected trough was paired with a subsequent peak enabling us to compute a peak to trough fold change ratio which is indicative of local signal amplitude. We then report the maximum peak to trough fold change per track, or the peak to trough fold change of the first peak of the track.

### Wavelet

To measure periods over time we used pyBOAT, a periodicity analyses platform ^57^. A cut-off period of 24 hours was used for sinc detrending, and the processed signal was then analysed using wavelet analysis ranging between 1 and 30 hours period. A Fourier estimate of the wavelet analysis provides a distribution of periods and their corresponding powers, and the period with the maximum power for each time point was considered. Then, those periods with powers above those found in background traces was used in the analysis.

### Analysis of single cell traces in R

To allow for comparison between datasets imaged on different microscopes/different days, the mVenus intensites were normalised to the maximum intensity of that experiment. Since cells did not start expressing NGN3::mVenus synchronously, all cell timeseries (tracks) were annotated with a “developmental time” and a “cell time”. Developmental time was in relation to the start of the population NGN3 window (day 15 of the differentiation), while cell time was in relation to the cell’s own NGN3::mVenus expression start time (when the Venus fluorescence became visible via imaging). The time in which NGN3 turns off per cell was calculated level-independently for each cell by calculating the last time in which the mVenus intensity reached the maximum intensity of that cell divided by 2.5.

Scarlet positive and scarlet negative cells from coculture differentiations were categorised using control experiments (positive and negative cell alone) and a bimodal mix model in R (mixtools package). For timeseries data, first, the background fluorescence of the experiment was removed from all the data points and the lower 5% quantile of the Scarlet intensity per cell was used for categorising to remove technical autofluorescent bright spots from the tracks. Any cell that was in a region of overlap between positive or negative controls, or in the overlap between the bimodal distributions was not taken forward for analysis.

The correlation matrix (Fig.S4A) was generated using the corrplot package in R, using hierarchical clustering (order = “hclust”) with two boxes drawn after seeing the results of the clustering (addrect = 2). Only correlation coefficients are shown if the p value > 0.001 (insig = “blank”.)

### RNA-Seq

RNA was extracted at days 0,3,6,9,12,15,18 and 24 of the differentiation using the WT FSPS13b hIPS cell line, with three biological replicates per time point. Unstranded, poly(A) selection, mRNA library preparation was performed and 75bp paired end sequencing, with approximately 40million reads per sample. Tophat v2 was used to align the reads to the reference human genome assembly (GRCh37/hg19) and the function ‘rpkm’ in the R/Bioconductor package edgeR was used with default parameters to normalize count gene expression.

### Bioinformatic analysis

Fig.6A heatmap was generated using pheatmap package in R with scaling by rows, complete clustering method, and Euclidean clustering distance measure. Genes were included if they are bound by NGN3, and if they are upregulated upon NGN3 expression, i.e. their expression on day 15 was higher than day 12 in the RNA-seq data.

A network of interactions from the ReMap 2022 database of published TF binding sites were used to assess whether cluster 1 genes (from Fig.6A) had binding sites associated with cluster 2 genes. All tissue types were included in this analysis.

### Incoherent Feed-Forward Loop

To provide a mathematical description of the resultant downstream effects of NGN3’s dynamics we put forward an incoherent feedforward loop (I1-FFL) motif. We developed a synthetic signal containing properties of NGN3 temporal dynamics as found in our data and provided this as an activatory input to two hypothesised downstream targets (X and Y where X also is a repressor of Y) and simulated the results.

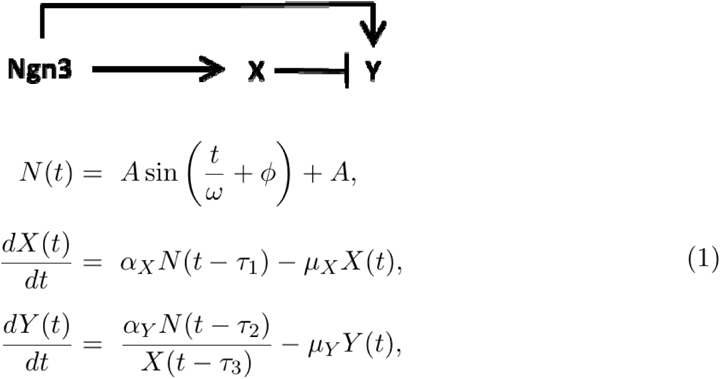

Where and Parameter represents half of the amplitude (peak to trough height), represents the periodicity (peak to peak temporal distance) and is the phase of oscillatory NGN3 dynamics, and denote the scalar constants representing the production and, the degradation of target genes and respectively. We set parameter values of, = 1 and = 0.1. Interactions between NGN3 and its targets X and Y are subject to time delays arbitrary time units (ABU). Additionally, there exists a time delay of ABUs for the repressive interaction from X onto Y. All parameter values were chosen as such to provide an accurate reflection of the network in question as the authors hypothesise it. To capture the properties of the gene expression dynamics, we prescribed our NGN3 simulations with a sinusoidal function where the amplitude and periodicity of each simulation within the same image varied by a given scale factor constant. Since the system of equations (1) include DDEs, we require history: these history functions of each consisted of vectors (with of length of)) full of constant values equal to. Furthermore, for Fig5.G, we terminated the sine wave in N(t) and enforced N(t) = 0 for all t after N reaches a trough after the first and third oscillatory peak in the red and blue lines respectively since these numbers of oscillatory peaks were frequently seen in NGN3s data (see Figure 1H). These troughs were found using the inbuilt “findpeaks” function in MATLAB. In all cases, the system of ODEs within (1) were solved using a fourth order Runge-Kutta numerical scheme in our MATLAB pipeline (found in the github repository). The solutions for the hypothesised target genes (X and Y) within the IFFL motif were computed along with the plot of a speculated “cumulative probability of differentiation” (in Figures 5H and J: bottom panel). Here we assumed that both X and Y target expression can affect the probability to differentiate in equal share. The cumulative probability was obtained by dividing the solution trace by the cumulative sum of each X and Y solution vector and then averaging the two. Peak to trough amplitude, peak to peak periodicity, and mean levels (A.U.) of the inputs along with initial conditions of components X and Y are given in Table S5. Note, the initial conditions of X were imposed depending on the mean levels of the NGN3 input as specified in Table S5.

## Data Availability

Raw imaging data is deposited at: https://figshare.com/projects/XXXXX

MATLAB pipeline for incoherent feed forward loop can be found in github (see repository https://github.com/Papalopulu-Lab/Miller2023)

RNA-seq raw data have been deposited at GEO and are publicly available at the time of publication at XXX.

## Notes

https://github.com/Papalopulu-Lab/Miller2023

### Competing Interest Statement

The authors have declared no competing interest.

