## Supplemental information for "NGN3 oscillatory expression controls the timing of human pancreatic endocrine differentiation"

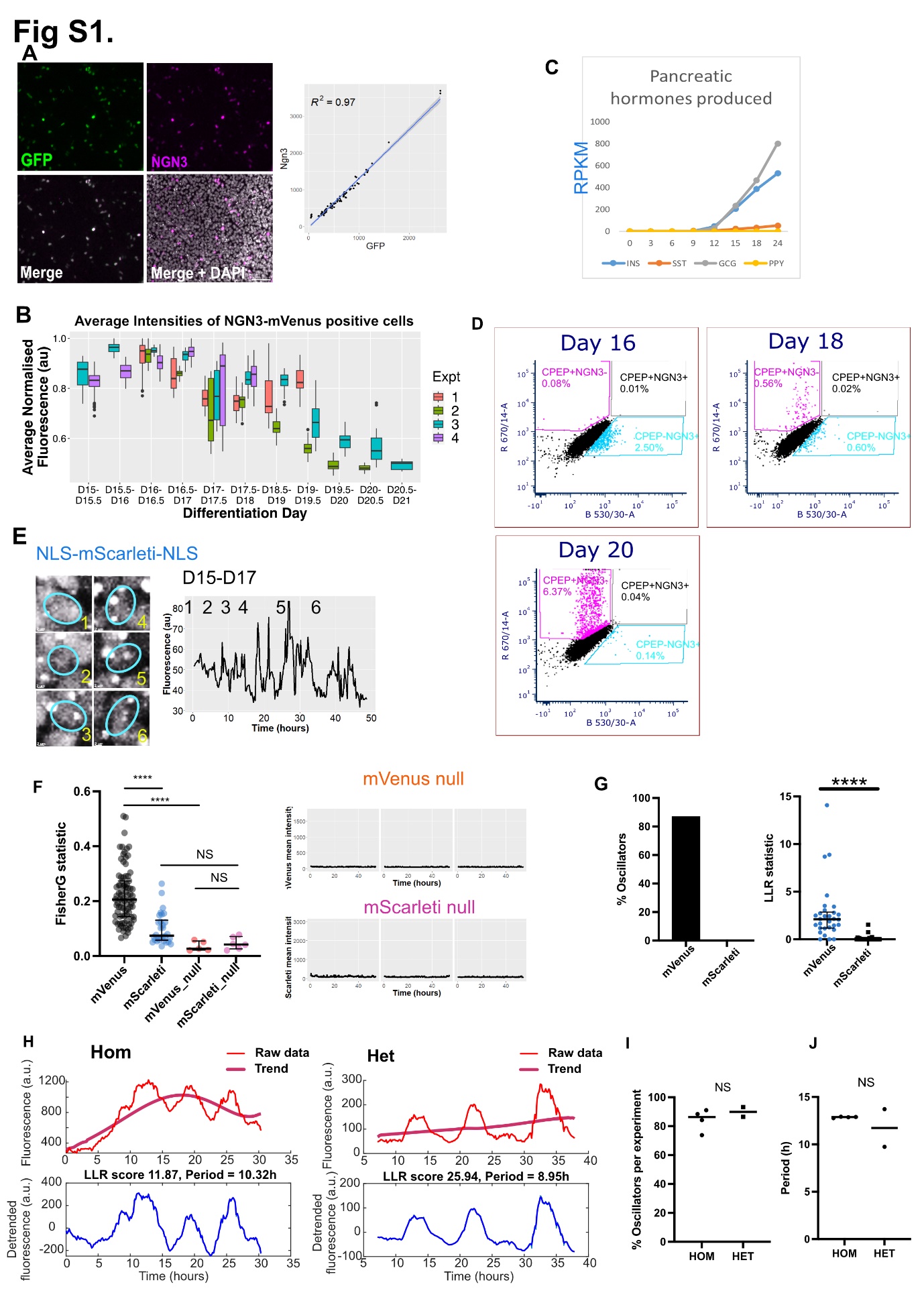

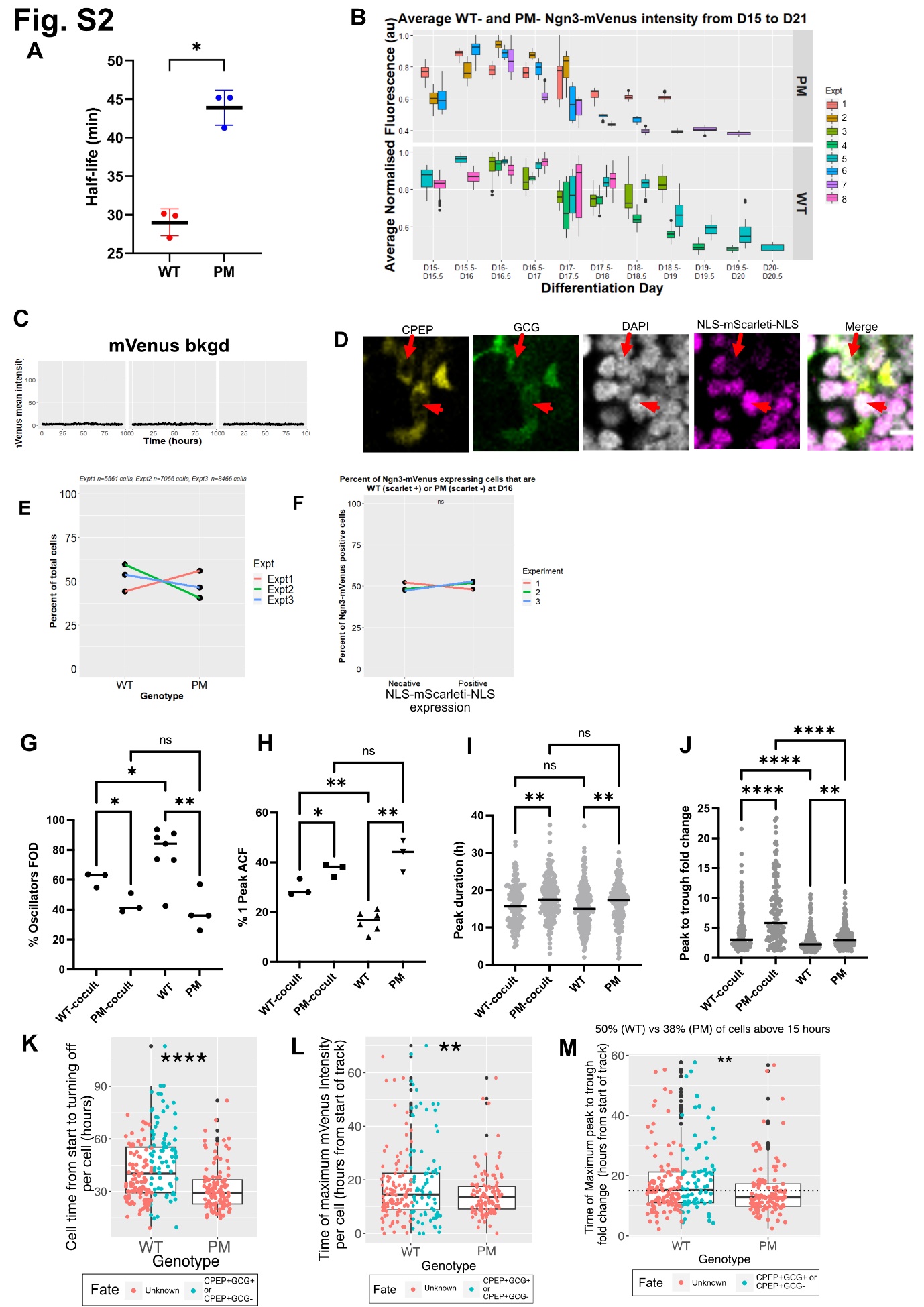

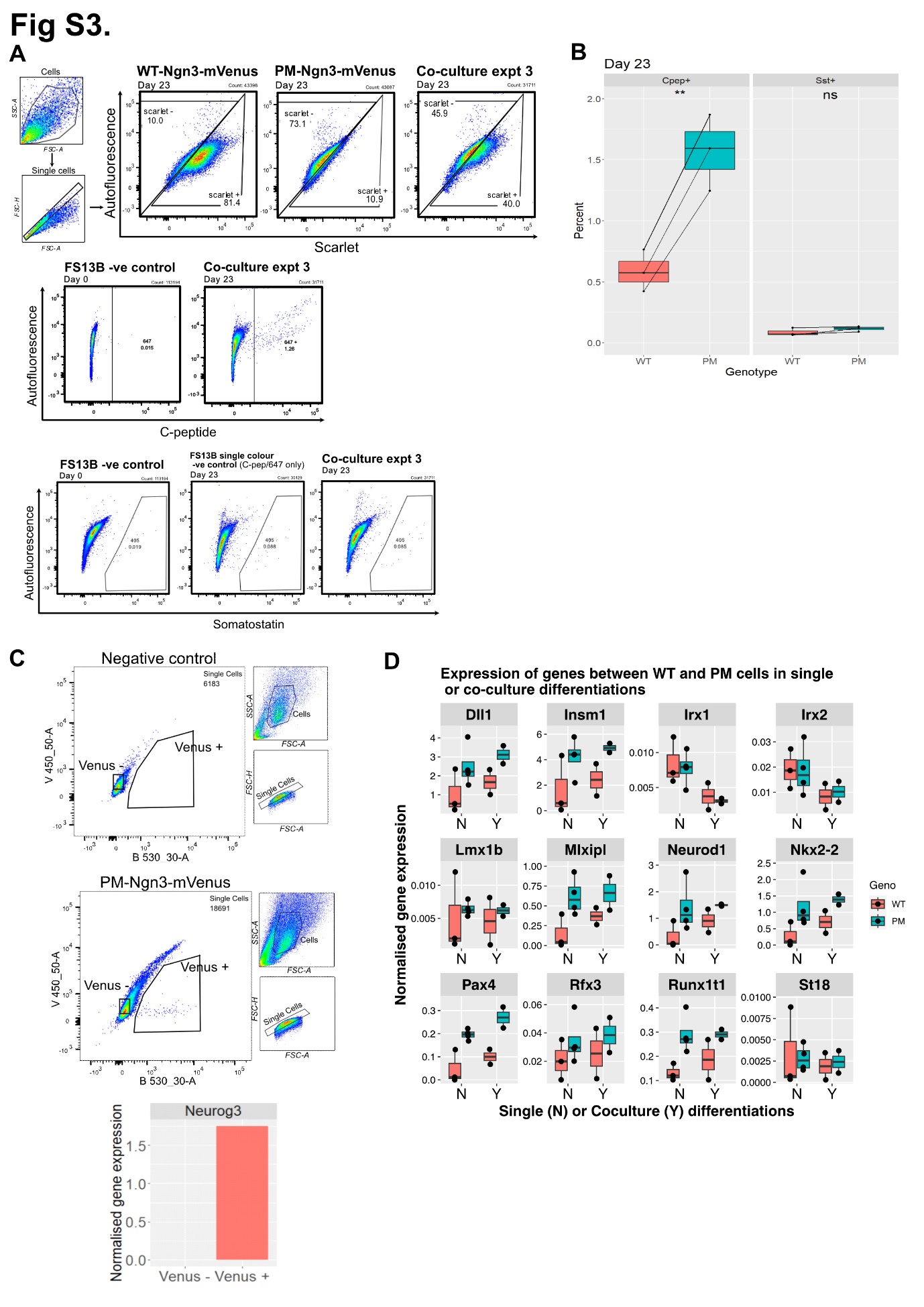

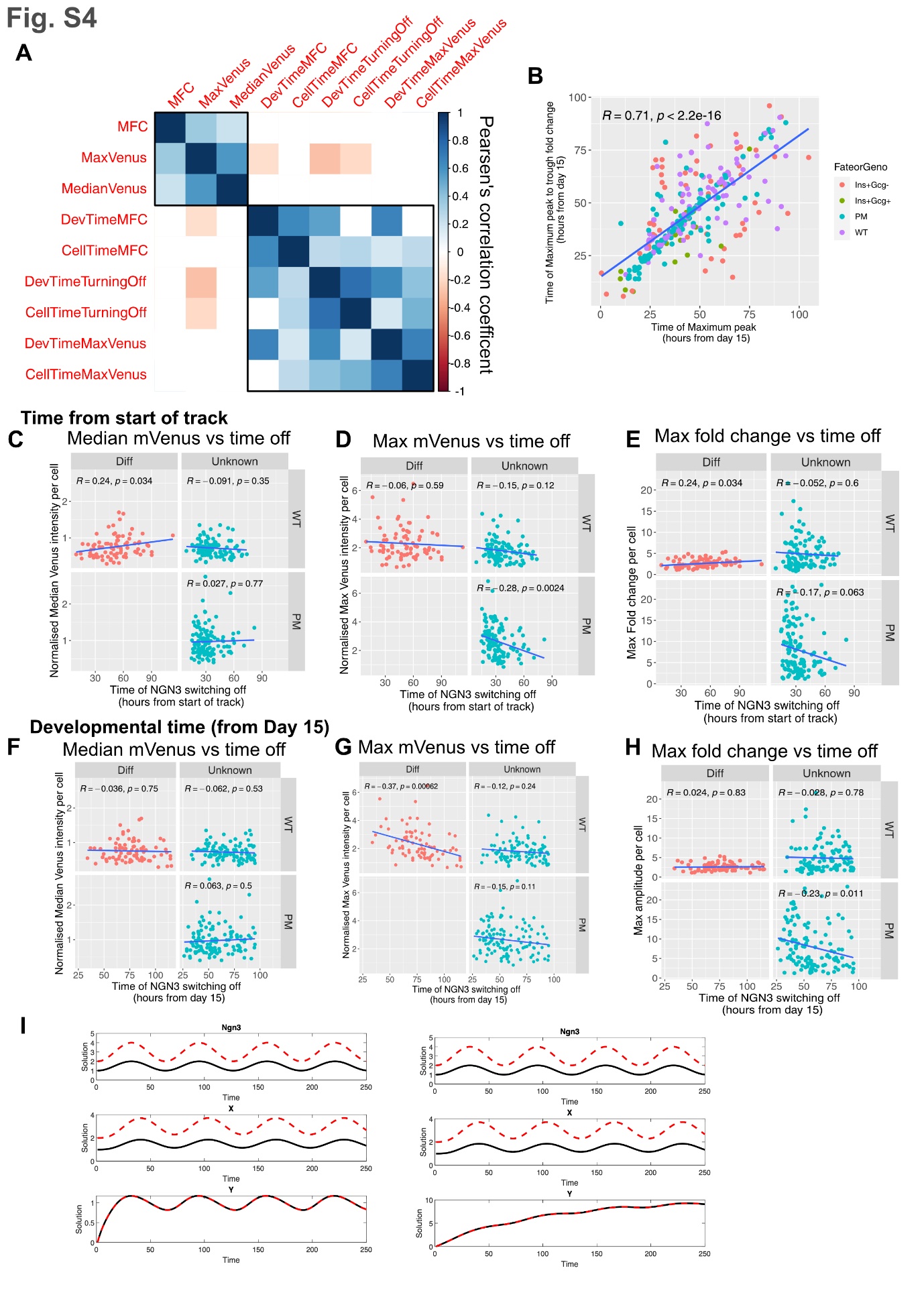

**Supplementary Figure Legends**

**Figures S1. Quantification of NGN3::mVenus and differentiation markers with experimental controls. Related to Fig 1.**

**(A)** Immunofluorescence of homozygous NGN3::mVenus+ cells (left panel) stained with α-NGN3 and α-GFP antibodies (detects Venus protein); quantification of GFP intensity per nucleus versus NGN3 intensity per nucleus indicates linear correlation with Pearson’s correlation coefficient R^2^ (right panel).

**(B)** Quantification of mVenus intensities of the positive cells from **(Fig.1D)** observed during pancreas differentiation days 15 to 20 across 4 independent experiments. The average intensity of each time point is normalised to the maximum intensity per experiment. Boxes indicate median and interquartile range; dots indicate outliers.

**(C)** Quantification of expression of pancreatic hormones INSULIN (INS), SOMATOSTATIN (SST), GLUCAGON (GCG) and PANCREATIC POLYPEPTIDE (PPY) over time from RNA-seq.

**(D)** Flow cytometry analysis showing detected fractions of CPEP+ in combination with NGN3+ during differentiation days 16 to 20.

**(E)** Timelapse images (left panel) and corresponding NLS-mScarleti-NLS expression (right panel-quantified as average intensity) observed in the nucleus in Fig 1G; images are numbered and nucleus is indicated with an elliptical boundary curve.

**(F)** Analysis of FisherG statistic from frequency spectra datasets (Materials and Methods) corresponding to average fluorescence fluctuations in NGN3:mVenus and NLS-mScarleti-NLS tracked in the same nuclei; high values indicate periodicity while low values indicate noise; the statistical value corresponding to detector noise is quantified from tracking nuclei of cells with no expression, referred to as mVenus_null and mScarlet_null; markers indicate nuclei, bars indicate median and interquartile range.

**(G)** Percentage of oscillatory NGN3:mVenus and corresponding NLS-mScarleti-NLS quantified in the same cells using a false discovery rate approach (left-panel, Materials and Methods); log-likelihood ratios (LLR) estimated from NGN3:mVenus and corresponding NLS-mScarleti-NLS signal observed in the same cells and quantified using a Gaussian Process approach (Materials and Methods), markers indicate nuclei, bars indicate median and interquartile range, statistical significance determined with a Wilcoxon matched-pairs signed rank test p<0.0001****.

**(H)** Representative examples of NGN3:mVenus periodic activity observed over time in single nuclei containing a homozygous (left panel) and heterozygous (right panel) endogenous knock-in; (top panels) de-trending of raw data; (bottom panels) detrended zero-mean data with Gaussian Model fit.

**(I-J)** Percentage of oscillators **(I)** and period estimates **(J)** compared in cells containing homozygoys (HOM) versus heterozygous (HET) NGN3:mVenus knock-ins; markers indicate median per experiment; line indicates median overall; unpaired t-test, two-tail non-significant (NS) p=0.4142**(I)** and p=0.4056**(J)**.

**Figure S2. Quantitative analysis of WT-NGN3 and PM-NGN3 in the same culture conditions. Related to Fig 3.**

**(A)** Quantification of half-life in cells hIPS expressing wild-type (WT) or PM versions of NGN3; dots indicate independent experiments; bars indicate mean and SD; statistical testing paired t-test, p=0.0186.

**(B)** Quantification of average WT-NGN3::mVenus and PM-NGN3::mVenus monitored in differentiation days 15 to 21 from data corresponding to **(Fig.3C)**; binning 12h, bars indicate median and interquartile range of individual experiments; dots indicate outliers. Venus intensities were normalised to the maximum intensity per experiment.

(**C)** Examples of mVenus-null expression monitored in NGN3- progenitors referred to as background.

**(D)** Representative single cell examples of multiple markers indicating C-peptide (CPEP), Glucagon (GCG), nuclear DAPI and nuclear NLS-mScarleti-NLS by immunofluorescence. Arrows indicate CPEP+GCG+mScarlet- cell, arrow head indicates a CPEP+GCG+mScarlet+ cell.

**(E)** Percentage of total WT versus PM cells at Day 16 in 3 independent co-culture experiments determined based on NLS-mScarleti-NLS marker expression (marking WT) using a Gaussian Mixture Model thresholding approach (Materials and Methods).

**(F)** Percentage of NGN3::mVenus positive WT and PM cells at differentiation day 16 determined based on NLS-mScarleti-NLS marker expression (marking WT) using the Gaussian Mixture Model thresholding approach from (E), normalised to the number of total number of cells in (E).

**(G-H)** Comparative analysis of percentage of oscillators **(G)** and percentage of tracks with one peak **(H)** observed mixed WT (WT-cocult) and PM (PM-cocult) conditions as well as unmixed WT and PM differentiation conditions; dots indicate independent experiments, lines indicate medians; statistical significance determined by one-way ANOVA followed with Fisher’s LSD test, p<0.05*, p<0.01**, p<0.0001*** and not-significant (ns-B, p=0.4944; ns-C, p=0.1025).

**(I-J)** Comparative analysis of peak duration **(I)** and maximum peak to trough fold change **(J)** observed in mixed WT (WT-cocult) and PM (PM-cocult) conditions as well as unmixed WT and PM differentiation conditions; gray markers indicate single cells, lines indicate medians; statistical significance determined by a one-way ANOVA followed by a Kruskal-Wallis test, p<0.05*, p<0.01** and p<0.0001**** and not-significant p=0.6050^ns1^ and p=0.1277^ns2^.

**Figure S3. Quantitative analysis of differentiation and NGN3 target gene expression in wild-type and phosphomutant NGN3 in separate and coculture differentiations. Related to Fig.4.**

**(A)** Flow cytometry analysis workflow including controls to determine the NLS-mScarleti-NLS, C-PEPTIDE and SOMATOSTATIN fractions to compare WT and PM cells in co-culture conditions at Day 23.

**(B)** Percentage of cells expressing C-PEPTIDE (CPEP+) and SOMATOSTATIN (SST+) observed in co-cultures of WT and PM from workflow shown in (A); dots and lines indicate paired experiments; bars indicate median; statistical significance determines paired t-test, p<0.01** and non-significant for p>0.05.

**(C)** FACS of mVenus expressing fractions of cells (top panels) and validation of NGN3 expression by RT-qPCR (bottom panel).

**(D)** Comparison of target gene expression in mVenus FACS sorted cells quantified by RT-qPCR at differentiation day 16 in coculture or independent WT and PM differentiation conditions; dots indicate independent repeats; bars indicate median and interquartile range.

**Figure S4. Correlation analysis of descriptors of NGN3 dynamics and mathematical exploration of target genes. Related to Fig. 5**

**(A)**Pearson’s correlation matrix clustered into two main groups, Max Fold Change (MFC), normalised maximum venus intensity per cell (Max Venus), normalised median venus intensity per cell (Median Venus), Time of the maximum fold change per cell in hours from day 15 off the differentiation (DevTime MFC) or from the start of the NGN3 expression per cell (CellTimeMFC), time the cell turns off NGN3 in hours from days 15 (DevTimeTurningOff) or from the start of the NGN3 expression per cell (CellTimeTurningOff), time off the maximum venus intensity per cell in hours from day 15 (DevTimeMaxVenus) or from the start of the NGN3 expression per cell (CellTimeMaxVenus). Colours represent the pearsen’s correlation coefficient, and only coefficients are shown if p value <0.001 (non significant squares are shown in white).

**(B)**Correlation analysis of the time of the maximum peak to trough fold change per cell versus the time of the maximum Venus intensity per cell observed in mixed WT and PM cultures, as well as WT cells that become CPEP+/GCG- and CPEP+/GCG+; in each panel, intensity values were normalised by the maximum of each experiment across 6 independent experiments; R values indicate Pearson’s correlation coefficient.

**(C)to(E)** Correlation analysis of the normalised median venus intensity per cell (C), normalised maximum venus intensity per cell (D) or the maximum peak to trough fold change per cell (E) versus the time of NGN3 turning off per cell in hours from the start of the NGN3 expression per cell . Each plot is split into WT and PM, as well as WT cells that become CPEP+/GCG- and CPEP+/GCG+ (grouped in Diff). Intensity values were normalised by the maximum of each experiment across 6 independent experiments (3 cocultures and 3 independent WT differentiations); R values indicate Pearson’s correlation coefficient.

**(F)to(H)** Correlation analysis of the normalised median venus intensity per cell (C), normalised maximum venus intensity per cell (D) or the maximum peak to trough fold change per cell (E) versus the time of NGN3 turning off per cell in hours from differentiation day 15. Each plot is split into WT and PM, as well as WT cells that become CPEP+/GCG- and CPEP+/GCG+ (grouped in Diff). Intensity values were normalised by the maximum of each experiment across 6 independent experiments (3 cocultures and 3 independent WT differentiations); R values indicate Pearson’s correlation coefficient.

**(I)** Simulations of the same model set up as Fig. 5E, but where the stability of X and Y are the same (left panel) or where Y is more stable than X (right panel).

Table S1

| **Antibodies** | **Company** | **Catalogue number** | **RRID number** | **Dilution used for IF** | **Dilution for Flow** |
| --- | --- | --- | --- | --- | --- |
| Anti-C-peptide | Origene | BM270S | AB_978884 | 1:500 | 1:100 |
| Anti-Glucagon | Abcam | ab92517 | AB_10561971 | 1:1000 | 1:100 |
| Anti-Somatostatin | Santa Cruz | Sc-47706 | AB_628268 | 1:100 | 1:50 |
| Anti-GFP | Roche | 11814460001 | AB_390913 | 1:500 | - |
| Anti-Ngn3 | R&D | AF3444 | AB_2149527 | 1:100 | 0.4µg |
| Donkey anti-sheep Alexa Fluor 568 | Thermo Fisher | A21099 | AB_2535753 | 1:1000 | 1:1000 |
| Donkey anti-rabbit Alexa Fluor 488 | Thermo Fisher | A21206 | AB_2535792 | 1:1000 | 1:1000 |
| Donkey anti-mouse Alexa Fluor 647 | Thermo Fisher | A31571 | AB_162542 | 1:1000 | 1:1000 |
| Donkey anti-rat dylight 405 | Stratech Scientific Ltd | 712-475-153-JIR | AB_2340681 | 1:800 | 1:800 |

Table S2

| **qPCR SYBR primers** | | |
| --- | --- | --- |
| Gene name | Forward sequence | Reverse sequence |
| *NEUROD1* | ATCAGCCCACTCTCGCTGTA | GCCCCAGGGTTATGAGACTAT |
| *DLL1* | CCAGGGTTGCACACTTTCTC | CTACTACGGAGAGGGCTGCT |
| *NEUROG3* | CGCTGCTCATCGCTCTCTA | CTCCGTCTCACGGGTCAC |
| *NKX2-2* | GGAGCTTGAGTCCTGAGGG | TCTACGACAGCAGCGACAAC |
| *PAX4* | TGCTGTGCAGAGATGATTCC | GCAAGAGAAGCTCAAGTGGG |
| *INSM1* | CAGGTGTTCCCCTGCAAGTA | CTTGTTGATGTGCCGCGTAA |
| *RUNX1T1* | TGCTTGGATGTTCTGAGTGC | CAGAGCTGCTTCTCGATGTG |
| *RPL7* | ACAAGCGTGGTTATGGCAAA | CTCATGAATCAAATCCTCCATGCA |
| *MLX1PL* | AGAACCGGCGTATCACACAC | gTGCTCACGAGCCCATGAA |

Table S3

| **qPCR Taqman Probes** | Product code from Life Technologies |
| --- | --- |
| *ACTB* | Hs01060665_g1 |
| *IRX1* | Hs00411782_m1 |
| *IRX2* | Hs01383002_m1 |
| *ST18* | Hs00608494_m1 |
| *LMX1B* | Hs00158750_m1 |
| *RFX3* | Hs01060440_m1 |

**Table S4**

| **Measurement** | **HOM (by experiment)** | **HET (by experiment)** | **PM (by experiment)** | **HOM Median** | **HET Median** | **PM**  **Median** |
| --- | --- | --- | --- | --- | --- | --- |
| Minimum track length | 21.83h; 21.67h; 23.17h; 23.75h. | 22.17h; 22.5h. | 21.83h; 23.75h; 27.75h; 21.83h. | 22.5h | 22.33h | 22.79h |
| Average track length | 42.48h; 27.95h;  36.91h; 40.99h. | 33.36h; 32.26h. | 29.7h; 37.47h;  53.56h; 26.53h. | 38.95h | 32.81h | 33.59h |
| Number of cells tracked | 86; 67; 80; 57. | 74; 37. | 43; 50; 138; 86. | 74 cells | 56 cells | 68 cells |
| Pearson’s Correlation Coefficient of track length vs LLR | (-0.1883;; -0.3463;; -0.1579;; 0.0338.) | 0.017;; 0.0867. | -0.3283;  0.0912;  -0.0268;  -0.1771. | -0.1731* | 0.05185* | -0.1020* |
| Pearson’s Correlation Coefficient of track length vs period | -0.1646; 0.064; 0.1248; 0.0059. | 0.0471; 0.1051. | 0.0778;  -0.1088;  -0.1269;  -0.0261. | 0.03495* | 0.07610* | -0.06745* |

* coefficients approx. 0 indicate no correlation

| Input signal | 2 x $\boldsymbol{A}$ (A.U.) | Mean of $\boldsymbol{N}$ (A.U.) | $\boldsymbol{\omega}$ (arbitrary time units) | $\boldsymbol{X(0)}$ | $\boldsymbol{Y(0)}$ |
| --- | --- | --- | --- | --- | --- |
| Fig 5 E black line | 1 | 1.5 | 63 | 1 | 0 |
| Fig 5 E red line | 2 | 3 | 63 | 2 | 0 |
| Fig 5 F black line | 10 | 6 | 63 | 1 | 0 |
| Fig 5 F red line | 2 | 6 | 63 | 1 | 0 |
| Fig 5 K red line | 2 | 1 | 63 | 1 | 0 |
| Fig 5 K blue line | 2.6 | 1.3 | 72.4 | 1 | 0 |
| Fig 5 N black line | 2, then 4, then 6 | 4 | 63 | 1 | 0 |
| Fig 5 N red line | 6, then 4, then 2 | 4 | 63 | 1 | 0 |

**Table S5**
